# Inference of Phylogenetic Networks from Sequence Data using Composite Likelihood

**DOI:** 10.1101/2022.11.14.516468

**Authors:** Sungsik Kong, David L. Swofford, Laura S. Kubatko

## Abstract

While phylogenies have been essential in understanding how species evolve, they do not adequately describe some evolutionary processes. For instance, hybridization, a common phenomenon where interbreeding between two species leads to formation of a new species, must be depicted by a phylogenetic network, a structure that modifies a phylogeny by allowing two branches to merge into one, resulting in reticulation. However, existing methods for estimating networks are computationally expensive as the dataset size and/or topological complexity increase. The lack of methods for scalable inference hampers phylogenetic networks from being widely used in practice, despite accumulating evidence that hybridization occurs frequently in nature. Here, we propose a novel method, PhyNEST (Phylogenetic Network Estimation using SiTe patterns), that estimates phylogenetic networks directly from sequence data. PhyNEST achieves computational efficiency by using composite likelihood as well as accuracy by using the full genomic data to incorporate all sources of variability, rather than first summarizing the data by estimating a set of gene trees, as is required by most of the existing methods. To efficiently search network space, we implement both hill-climbing and simulated annealing algorithms. Simulation studies show that PhyNEST can accurately estimate parameters given the true network topology and that it has comparable accuracy to two popular methods that use composite likelihood and a set of gene trees as input, implemented in SNaQ and PhyloNet. For datasets with a large number of loci, PhyNEST is more efficient than SNaQ and PhyloNet when considering the time required for gene tree estimation. We applied PhyNEST to reconstruct the evolutionary relationships among *Heliconius* butterflies and Papionini primates, characterized by hybrid speciation and widespread introgression, respectively. PhyNEST is implemented in an open-source Julia package and publicly available at https://github.com/sungsik-kong/PhyNEST.jl.

## 1 Introduction

The relationships among organisms are often portrayed in a tree-like structure called a phylogenetic tree or phylogeny. Phylogenetic trees assume that the formation of a novel lineage occurs in a bifurcating manner where an ancestor splits into two descendants that possess traits vertically inherited from their common ancestor. This structural constraint overlooks important and common biological phenomena such as hybridization, an umbrella term that includes both hybrid speciation and introgression (Anderson, 1953), that can also result in the production of a new lineage (Patterson et al., 2012; Ungerer et al., 1998; Grant and Grant, 1992; Mallet, 2007; Mavárez et al., 2006; Rieseberg and Carney, 1998; Abbott et al., 2013; Arnold, 1992; Lamichhaney et al., 2018). In this case, genetic material is transferred horizontally between organisms, and a phylogenetic network, a graph that generalizes a phylogeny by allowing two branches to merge into one at a node and create a structure called a reticulation, is a more accurate representation of the true evolutionary history (Huson et al., 2010; Huson and Bryant, 2006; Blair and Ané, 2020; Kong, 2022).

While the term phylogenetic network is often used to refer both abstract and explicit networks in the literature, only explicit networks depict evolutionary history (Morrison, 2005; Kong et al., 2022). In fact, abstract networks are unsuitable for evolutionary investigations because they cluster sequences based on overall similarity of their nucleotide arrangement, lack evolutionary directionality, and contain cycles that do not represent complex evolutionary histories but rather conflicting signals in the data (Kong et al., 2016; Sánchez-Pacheco et al., 2020). In contrast, explicit networks are often rooted and directed, and statistically inferred. Explicit networks also include recently introduced unrooted, semi-directed phylogenetic networks (Solís-Lemus and Ané, 2016; Allman et al., 2019; Xu and Ané, 2021) where the directionality is eliminated except for the reticulation edges. Hereafter, we refer to explicit networks when we say phylogenetic networks (or networks).

Significant developments in phylogenetic network inference methods from multi-locus data have been made, but scalability remains a major challenge since the computational cost becomes large as the dataset size (i.e., number of individuals and/or sequence length) and/or topological complexity (i.e., number of reticulations) increase. One common approach to alleviate the computational burden is to summarize an input sequence alignment through estimation of gene trees for each locus prior to network estimation (e.g., PhyloNet (Than et al., 2008; Wen and Nakhleh, 2018), SNaQ (Solís-Lemus and Ané, 2016), NANUQ (Allman et al., 2019), RF-NET2 (Markin et al., 2022)). While data summarization improves efficiency, inference of networks from the gene trees is still computationally intensive (Hejase and Liu, 2016). Such methods are also prone to gene tree estimation error (Yu et al., 2014), although counterstrategies exist like use of concordance factor (CF) in SNaQ and a set of gene trees per locus as input in PhyloNet. Most importantly, the summarization can result in the loss of relevant phylogenetic signal in the data. While some methods can estimate networks directly from an input alignment (e.g., BEAST 2 (Zhang et al., 2018), BPP (Flouri et al., 2020), the Bayesian methods in PhyloNet (Wen and Nakhleh, 2018)), they are impractical for large datasets as their computation involves simultaneous inference of topologies for both genes and species.

Here, we propose an efficient and statistically-sound method to estimate phylogenetic networks without the need to summarize an input sequence alignment into a set of gene trees. Our method is based on a composite likelihood framework and is called PhyNEST (Phylogenetic Network Estimation using SiTe patterns). In the following sections, we formally introduce relevant models and provide a detailed description of the network inference procedure. Using simulation, we assess the efficiency and accuracy of PhyNEST and evaluate its performance in comparison to two popular summary-based methods that use composite likelihood implemented in PhyloNet and SNaQ. Finally, we apply PhyNEST to estimate evolutionary relationships among *Heliconius* butterflies (Martin et al., 2013), where evidence of hybridization is strong in previous studies (Mavárez et al., 2006; Brower, 2013), and Papionini primates, where rampant introgression was reported (Vanderpool et al., 2020). PhyNEST is implemented in Julia programming language and publicly available at https://github.com/sungsik-kong/PhyNEST.jl.

## 2 Model

### 2.1 Trees and Networks

#### 2.1.1 Phylogenetic trees

Let *X* = {*x*_1_, *x*_2_,…, *x_n_*} denote a non-empty finite set of *n* taxa. In a phylogenetic tree (Fig. 1a and 1b), the genealogical relationships among the elements in *X* are primarily represented in the form of topology *T*, a tree with no branch length specified. A topology consists of a set of nodes *V* (*T*) and a set of branches *E*(*T*), so we define *T* = (*V* (*T*), *E*(*T*)).

**Figure 1:**
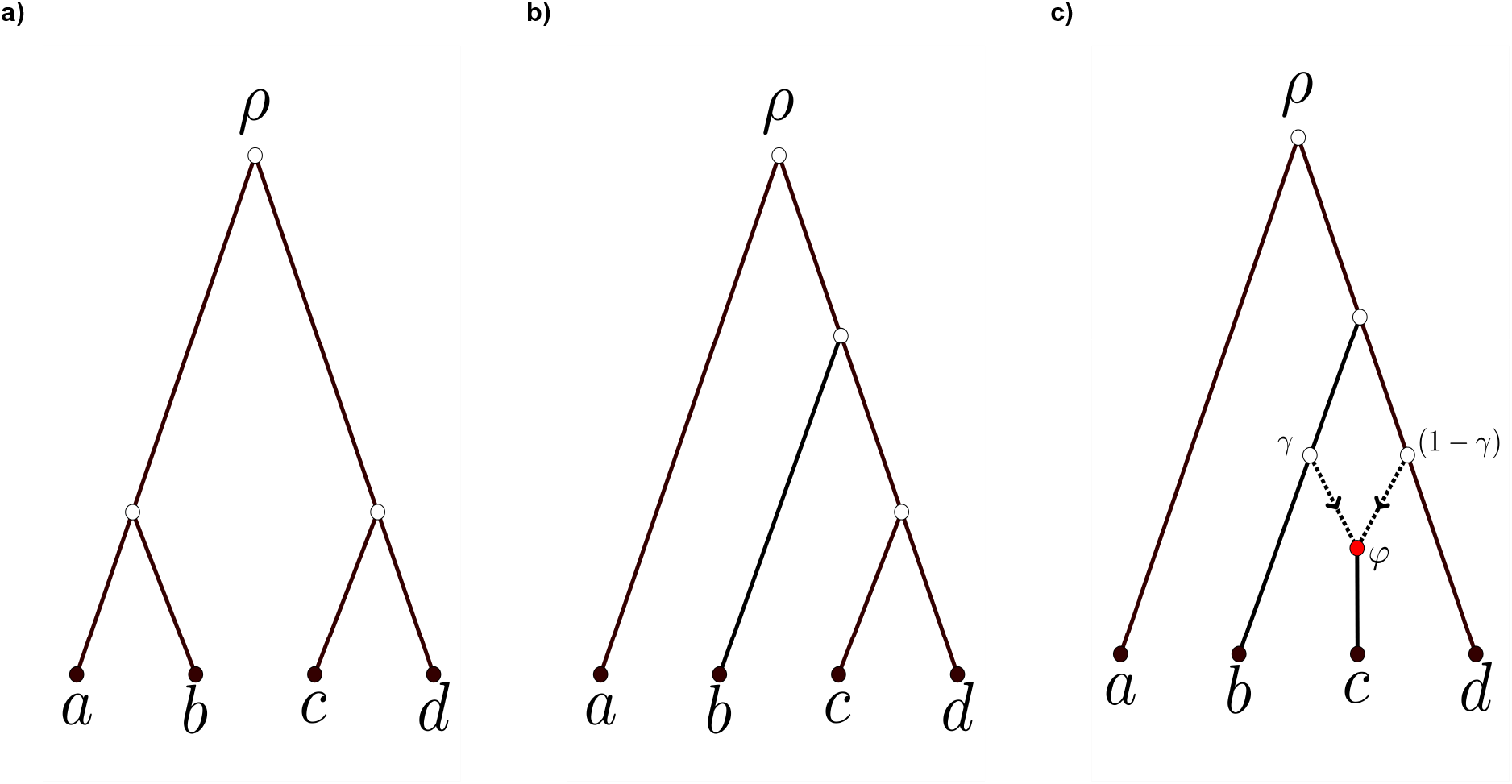
Examples of ((a) and (b)) phylogenetic trees and (c) a network and on *X* = {*a, b, c, d*}. All edges are directed down the page beginning from the root *ρ* that represents the most recent common ancestor of the taxa in *X*. The root and internal tree nodes are shown in hollow circles that represent speciation events. The leaves are shown in filled black circles that are bijectively labeled with *X* at the bottom of the tree or network. (a) and (b) represent symmetric 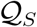 and asymmetric 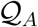 quartets, respectively. In (c), reticulation vertex *ϕ* is represented with a filled red circle that has two incoming reticulation edges (dotted and directed), with inheritance probability *γ* for the edge that connects to the ancestor of *b* and (1 − *γ*) for the edge that connects to the ancestor of *d*. The reticulation vertex *ϕ* represents a hybridization event between the ancestors of *b* and *d*, that resulted in hybrid taxon *c*.

The set of nodes *V* (*T*) = *ρ* ∪ *V_I_* ∪ *V_L_* satisfies the following properties:

i. *ρ* is the (unique) root of *T* that has in-degree zero and out-degree two;
ii. *V_I_* is the set of internal tree nodes that have in-degree one and out-degree two;
iii. *V_L_* is the set of leaves (or external tree nodes) that have in-degree one and out-degree zero.

The root *ρ* represents the most recent common ancestor of all sampled *V_L_* and inferred *V_I_*, and transforms the branching graph into a biological hypothesis by providing the evolutionary direction from *ρ* outward to *V_L_*. The leaves are bijectively labeled by the leaf-labeling function *f*: *V_L_* → *X* (i.e., each element of *V_L_* is identified with an element of *X*). Each element of *V_I_* represents an ancestral species (and a speciation event). A polytomy, a topology with one or more *V_I_* (or *ρ*) with out-degree > 2, is not included in our definition.

We denote the set of branches as *E*(*T*) ⊆ *V* (*T*) × *V* (*T*), and each branch as *e* = (*u, v*), where *u* is a tail (or a parent) of *v* and *v* is a head (or a child) of *u*. If a directed path between *u* and *v* exists in *T*, we interpret that *u* is an ancestor of *v*, and *v* is a descendant of *u*. Every branch in a phylogeny is a tree branch whose head is either in *V_I_* or *V_L_* and *e* ∈ *E*(*T*).

A metric tree refers to a topology with a collection of non-negative branch lengths specified that incorporates temporal information into the topology. Let *τ* be a vector of speciation times (synonymous to node ages) with *τ_i_* ∈ (0, ∞) for *i* = 1, 2,… *n* − 1 where *τ_i_* represents the time of the *i*th speciation from the present. Therefore, a metric phylogenetic tree comprised of both *T* and *τ* can be written as 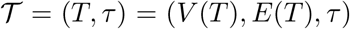. When every element of *V_L_* in 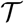 is equidistant from *ρ*, we say 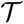 is ultrametric and satisfies the molecular clock, assuming that the mutation rate *μ* is constant throughout the entire tree.

#### 2.1.2 Phylogenetic networks

The topology of a phylogenetic network *N* (Fig. 1c) contains a set of vertices (synonymous to nodes in trees) *V* (*N*) = *ρ* ∪ *V_I_* ∪ *V_L_* ∪ *V_N_* where each element shares the same definition as in *V* (*T*) except that reticulation vertices in *V_N_* must satisfy the following property:

(iv) *V_N_* is the set of reticulation vertices that have in-degree two and out-degree one.

We denote *h* as the number of reticulation vertices. We exclude the case where a reticulation vertex has in-degree three or more, assuming it is biologically infeasible to have more than two parents.

The set of edges (synonymous to branches in trees) in *N*, *E*(*N*), is partitioned into tree edges (see *E*(*T*) above) and reticulation edges whose head is a reticulation vertex. Let *γ* represents a vector of inheritance probabilities in *N*, where an inheritance probability, denoted by *γ_j_* ∈ [0, 1], *j* = 1, 2,…, *h*, is associated with one of the two incoming reticulation edges for each reticulation vertex. In brief, *γ_j_* describes the proportion of the genome that the taxon represented by the corresponding reticulation vertex inherited from one of its parents, which in turn makes the contribution from the other parent as (1 − *γ_j_*) (Meng and Kubatko, 2009; Yu et al., 2012).

A phylogenetic network 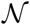 on *X* with *h* reticulations is defined as 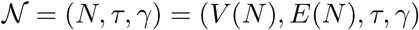 Note that *τ* does not include times of reticulation events and 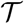 is one case of 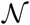 where *h* = 0. The networks that PhyNEST estimates are time-consistent tree-child (TCTC) level-1 networks as these networks make biological sense and are computationally tractable (see Kong et al. (2022) for a formal definition). The biological motivation behind time-consistency is that two taxa involved in a hybridization event must have temporally coexisted in order to interbreed (Cardona et al., 2014). In theory, the two incoming edges should have an edge length equal to zero since hybridization is an instantaneous process. However in practice, the lengths of reticulation edges can be larger than zero in cases of incomplete sampling or the presence of extinct taxa.

The reticulations in a level-1 network do not overlap (i.e., do not share an edge), thus restricting the level of a network to one leads to more computationally tractable problems, particularly in terms of statistical identifiability. More importantly, a level-1 network can be decomposed into a collection of parental trees (or displayed trees) that is obtained by eliminating any reticulation in the network through the removal of one of the reticulation edges for each reticulation node (Döcker et al., 2019). A network with *h* reticulations displays 2^*h*^ parental trees and we denote a parental tree of 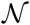 as PT^*t*^ where *t* = 1,…, 2^*h*^. A network is called a tree-child network if every non-leaf vertex is the parent of a tree vertex or a leaf. Biologically, this means that the populations that are involved in hybridization event always have some descendants that did not participate in the hybridization event but rather evolve solely through mutation (Kong et al., 2022).

### 2.2 Site Pattern Probability Distributions

The coalescent models the evolutionary history of a sample of lineages within a population backward in time (i.e., from a leaf towards the root), and the extension of the coalescent to the case of multiple taxa whose evolutionary history is represented by a species tree is called the multispecies coalescent (MSC; Kingman, 1982, 2000; Kubatko, 2019; Liu et al., 2009; Takahata, 1989). Occasionally, the genealogical history of the sampled populations for a particular locus, known as the gene tree, conflicts with the species tree due to processes like incomplete lineage sorting (ILS) and hybridization (Maddison, 1997; Pollard et al., 2006; Morales-Briones et al., 2021). Phylogenomic approaches based on MSC take the gene tree–species tree conflict caused by ILS into account (Mirarab et al., 2014; Liu et al., 2010; Chifman and Kubatko, 2014, 2015; Larget et al., 2010; Ane et al., 2007). Moreover, the recently proposed network multispecies coalescent extends these ideas by simultaneously modeling hybridization and ILS to infer the species-level relationships in a phylogenetic network (Degnan, 2018). Using the MSC to compute the probability distribution of gene trees given a particular species tree and set of speciation times (Degnan and Salter, 2005; Rannala and Yang, 2003), Chifman and Kubatko (2014, 2015) showed that it is possible to obtain analytic expressions for the probability distribution of data patterns (i.e., site patterns) at the tips of a species tree with four taxa. In this section, we describe the concept of a site pattern probability, provide details of the computation of site pattern probabilities given the relevant parameters on a quartet, and introduce the computation of site pattern probabilities on a quartet with reticulation.

#### 2.2.1 Site pattern probabilities on a quartet tree

Define a site pattern arising from tree 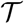 with *n* tips as an assignment of nucleotide states *s*_1_*s*_2_ … *s_n_* where *s_y_* ∈ {*A, C, G, T*}, *y* = 1, 2,…, *n*, to the tips of the tree. We assume each site is an independent observation from the species tree under the MSC, a data type referred to as coalescent independent sites (CIS). Note that there are 4^*n*^ possible site patterns in a tree with *n* tips. Coalescent independent sites refer to columns in a sequence alignment where all nucleotides evolved from a common ancestor according to some evolutionary process (Tian and Kubatko, 2017), and should not be confused with single nucleotide polymorphism data, as CIS data include constant sites as well (i.e., sites for which all taxa have the same nucleotide state).

Define a quartet tree 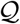 as a tree with four tips where one lineage is sampled per tip (e.g., Fig. 1a and 1b), thus 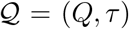 where *Q* and *τ* represent the topology and the vector of three speciation times of the particular 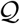, respectively. Let Y_*w*_ be the observed nucleotide at a site in the data at the tip *w* = 1, 2, 3, 4 in 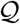. The probability of observing site pattern of *s*_1_*s*_2_*s*_3_*s*_4_ is defined as:

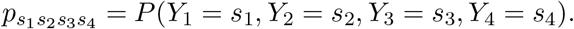

A split refers to a bipartition of *X* into two non-overlapping subsets of *X*, *X*_1_ and *X*_2_, and is denoted by *X*_1_|*X*_2_. A split is valid for 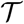 if the subtrees containing the taxa in *X*_1_ and *X*_2_ do not intersect. In case of 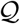, we consider splitting the four taxa into two groups of two (i.e., |*X*_1_| = 2 and |*X*_2_| = 2). Under this partition, the probability distribution 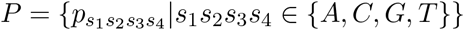 can be displayed in the form of a flattening matrix as shown in Chifman and Kubatko (2014) as:

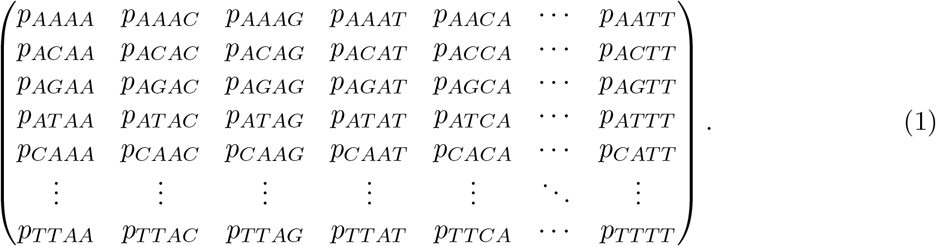

The 16 × 16 matrix above summarizes the 4^4^ = 256 possible site patterns for 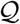. The rows and columns represent the possible nucleotide for the two taxa in the sets *X*_1_ and *X*_2_, respectively. These patterns can be reduced to 15 unique site pattern probabilities under the Jukes–Cantor model (JC69; Jukes and Cantor, 1969):

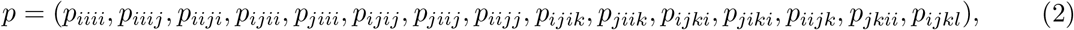

where *i*, *j*, *k*, and *l* denote distinct nucleotide states. We define the weight (*ω*) for each element in (2) as a vector of integers where each integer represents the number of occurrences of that site pattern in (1), in the same order as in (2):

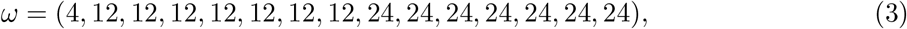

and 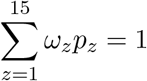 for *z* = 1, 2, …, 15.

#### 2.2.2 Computing site pattern probabilities on quartet trees and networks

Under the JC69 model, some of the 15 site pattern probabilities in (2) have equal probabilities (Peng et al., 2022). In the following, we show that the site pattern probabilities for 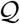 can be written as a function of the effective population size parameter *θ* and *τ* (Chifman and Kubatko, 2015).

Let 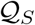 (Fig. 1a) and 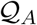 (Fig. 1b) denote the symmetric and asymmetric rooted quartets, respectively. In the case of 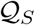, the 15 site pattern probabilities in (2) can be reduced to nine distinct probabilities as:

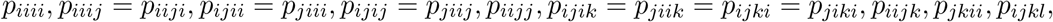

where any one of these site pattern probabilities can be computed with:

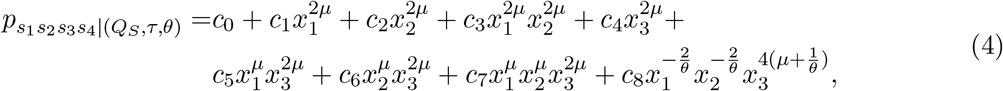

when 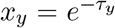 for *y* = 1, 2, 3, *μ* = 4/3, and the coefficients *c_w_* for *w* = 0, 1, 2, …, 8 are as shown in Table 1 in Appendix A. Let (*C_S_*)_9×9_ be the matrix of the coefficients and

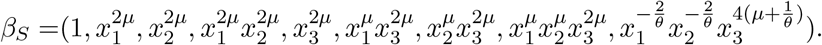

Then (4) can be written in matrix form as

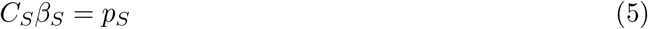

where

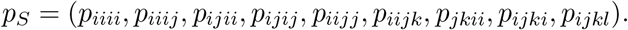

Similarly, in the case of 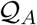, the 15 site pattern probabilities in (2) are reduced to 11 distinct site pattern probabilities as:

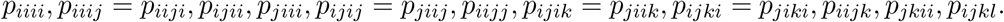

Each site pattern probability can be computed with:

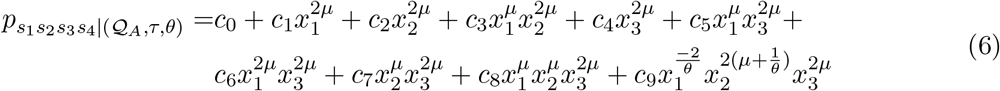

where 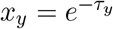 for *y* = 1, 2, 3, *μ* = 4/3, and the coefficients *c_z_* for *z* = 0, 1, 2, …, 9 for each of the 11 site patterns are given in Table 2 in Appendix A. Let (*C_A_*)_11×10_ be the matrix of the coefficients and

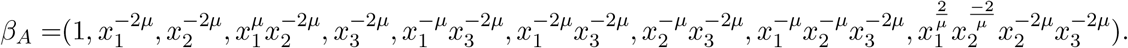

Then (6) can be written in a matrix form as

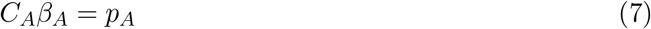

where

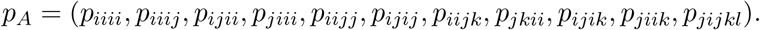

Define a quartet network 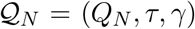 where *Q_N_*, *τ*, and *γ* represent a network topology with four tips, the vector of speciation times, and the vector of inheritance probabilities, respectively. Recall that a network 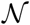 with *h* reticulations can be decomposed into a collection of 2^*h*^ parental trees, and any parental tree obtained from 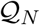 is either 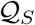 or 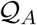. Let Γ represent the vector of the weights of the parental trees where Γ_*t*_, *t* = 1, 2, …, 2^*h*^, is the product of inheritance probabilities of all reticulation edges that are not removed in the production of *PT^t^*. Note that the sum of the elements of Γ must be 1. The site pattern probabilities for 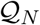 are the weighted sum of the site pattern probabilities of each *PT^t^* for that subset of four taxa, or 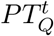:

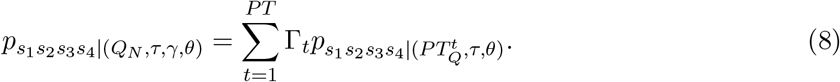

While the 15 site pattern probabilities on a quartet can be reduced to nine and 11 in the cases of 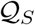 and 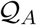, respectively, they are not reduced for 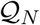 since the quartets extracted over the parental trees of 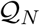 can be a mixture of 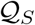 and 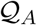, which leads to:

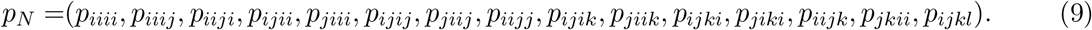

### 2.3 Computing the Composite Likelihood of a Network

Composite likelihood (or pseudolikelihood) refers to the formation of a likelihood function via multiplication of components for subsets of the data, even though the subsets may not be independent from one another (Varin et al., 2011). A composite likelihood function using site pattern frequencies for a phylogenetic tree was proposed in Peng et al. (2022) by decomposing an *n*-taxon tree into a set of quartets that includes all possible choices of four taxa. In brief, the likelihood of a quartet tree given a set of observed site pattern probabilities can be written using the site pattern probabilities in (4) and (6). The product of these quartet likelihoods is then used to compute the composite likelihood for the entire tree. In this subsection, we extend this idea to the network setting.

There are 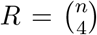 quartets in a network where *n* represents the number of leaves. For each site *m* in an alignment with length *M*, and for each quartet *q*, *q* = 1, 2,…, *R*, we define 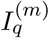 to be the random vector of length 15 that contains a 1 in the *j*th entry that corresponds to one of the 15 site patterns in (9) if that site pattern is observed at site *m* and 0 in all other entries. Let 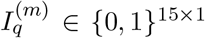 represent the corresponding observed data. Let *W_q_* be the vector of length 15 that counts of number of times the site patterns are observed. The *j*th component of *W_q_*, (*W_q_*)_*j*_, that represents the frequency of one of the site patterns is denoted by:

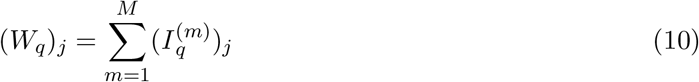

when 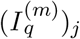 is the *j*th entry of 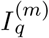.

Given the vector of observed site pattern frequencies *W_q_*, the likelihood for a quartet *ℓ_q_* is expressed as a function of *τ*, *θ*, and *γ* as:

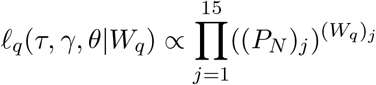

where (*P_N_*)_*j*_ is the *j*th entry in *P_N_* for quartet *q* and (*W_q_*)_*j*_ represents the *j*th entry of the vector for *q* that counts of number of times site pattern *j* is observed. Finally, we optimize the function

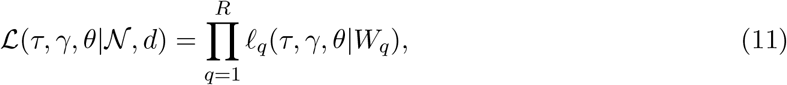

to compute the composite likelihood of 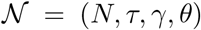 given the sequence data matrix *d* = (*I*^(1)^, *I*^(2)^,…, *I*^(*M*)^) where *I*^(*m*)^ is data at site *m* for the entire tree. The dimension of *d* depends on *n* and *M*. More specifically, each vector *I*^(*m*^ records which of the possible distinct site patterns on a tree of *n* tips is observed at site *m*.

## 3 Network Inference

### 3.1 Input Data

PhyNEST requires two inputs: (1) a multi-locus sequence alignment and (2) a starting topology. The sequence alignment refers to a plain text file containing DNA sequence data in sequential PHYLIP format. The starting topology is a Newick-formatted tree or an extended Newick-formatted network (Cardona et al., 2008), with or without *τ* and *γ* specified, and can be randomly generated or estimated from the data. PhyNEST uses the function `readTopology` in the Julia package PhyloNetworks (Solís-Lemus et al., 2017) to read in the input topology.

The site pattern frequencies for every unique combination of four sequences are computed followed by the rearrangement the site pattern frequencies for all permutations of four taxa. For each pair consisting of a quartet and an array of the corresponding site pattern frequencies, the rearrangement will results in 4! new combinations of taxa within a quartet, with a corresponding set of site pattern frequencies (see Table 3 in Appendix B). PhyNEST conducts checkpointing at the end of data parsing and creates a `.ckp` file that can be used as input in subsequent analyse, bypassing redundant treatment of the same dataset. Moreover, PhyNEST has an option to save all computed site pattern frequencies in a `.csv` file.

### 3.2 Optimizing the Objective Function

Optimizing the composite likelihood over the parameter set for a fixed topology requires finding the values of *τ, γ*, and *θ* that maximize the objective function (11). Optimization in PhyNEST uses the Broyden–Fletcher–Goldfarb–Shanno (BFGS) algorithm (Fletcher, 2000) implemented in Optim.jl, a Julia package for function optimization. Since the BFGS algorithm minimizes the objective function, the negative log of the objective function is optimized. In this section, we describe the parametrization of *τ*, *γ*, and *θ* used to transform the parameters to allow efficient unconstrained optimization, as well as determination of reasonable starting points for *τ* and *θ* to reduce the number of function evaluations needed for optimization. We expand ideas presented in implementation of qAge in PAUP* (Peng et al., 2022) to the network setting. These strategies are expected to enhance the efficiency of the optimization process overall.

#### 3.2.1 Parametrization and transformations

Consider a network with *n* tips with *h* reticulations where each internal tree vertex is indexed by an integer *j* = 1, 2,…, *J*, *J* = *n* + *h* − 1. The order of the labels is given by a postorder traversal with vertex age *τ_j_* for a vertex with index *j*. Let *A*(*j*) denote the index of the most recent ancestor of a vertex *j* (i.e., the first vertex that appears from *j* towards the root *ρ*) and let *ρ* have index *J*. Each reticulation vertex is indexed by an integer *r* = 1, 2,…, *h* where the inheritance probability *γ_r_* is given to one of the two incoming edges for the reticulation vertex. The other edge has inheritance probability 1 − *γ_r_*.

Numerical optimization of (11) is a nonlinear optimization problem that must satisfy the constraints that *τ*, *γ*, and *θ* are nonnegative, that *τ_j_* < *τ*_*A*(*j*)_, and that *γ_r_* ∈ [0, 1]. The free parameters being optimized are *τ*_1_, *τ*_2_,…, *τ_J_*; *γ*_1_,…, *γ_h_*; and *θ*. Note when *h* = 0 (i.e., a tree), *γ* is not a parameter to be optimized. In order to recast the inequality constrains *τ_j_* < *τ*_*A*(*j*)_ to simple bound constraints, we reparamtrize vertex ages using age ratios *τ_j_*/*τ*_*A*(*j*)_ for vertices 1, 2,…, *J* − 1, which are restricted to the interval [0,1]. Letting *i* = 1, 2,…, *J* + *r* +1, entries in the full parameter vector *η* are then defined as:

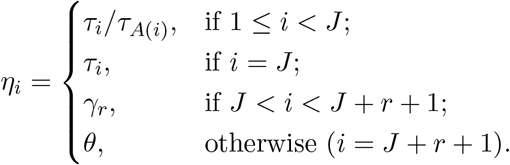

For each point *η* being evaluated during the optimization, we convert the *η_i_* corresponding to non-root vertex ages (i.e. *τ_i_* when *i* ≠ *J*) back to *τ_i_* prior to evaluating the composite likelihood function by:

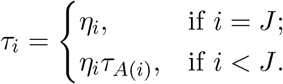

Note that *τ*_*A*(*i*)_ is always updated prior to using it for calculation of *τ_i_*. The optimization procedure must support simple lower and upper bounds on the variables in order to satisfy the constraints:

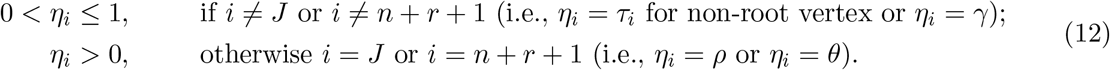

We converted the optimization problem to a fully unconstrained one by eliminating the bound constraints in (12) using suitable transformations (Box, 1966; Sisser, 1981). Arcsine square root transformations are used for *τ_i_*/*τ*_*A*(*i*)_ and *γ*; log transformations are used to enforce nonnegativity of the root age (= *τ_J_*) and *θ*. After these transformations, the parameter vector *ζ* becomes:

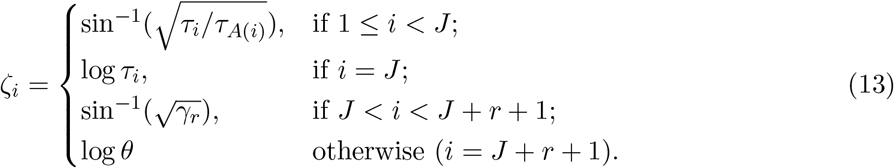

The parameter values in *ζ* are backtransformed to evaluate (11) using:

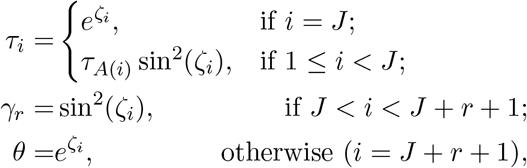

where the vertices indexed by *i* are backtransformed in preorder so that *τ*_*A*(*i*)_ is updated prior to using it for calculation of *τ_i_*.

#### 3.2.2 Starting values for *τ* and *θ*

Initiating the optimization procedure with a reasonably selected set of starting values of parameters is expected to enhance computational efficiency by reducing the number of function evaluations required. For optimization of the composite likelihood function (11), we can compute starting values for *τ* and *θ* given the site pattern frequencies acquired from the data for each quartet present in a network. In this section, we first describe obtaining starting values for *τ* when *θ* is known followed by the description of obtaining starting values for both *τ* and *θ* implemented in PhyNEST.

The ages of the three internal nodes *τ*_1_, *τ*_2_, and *τ*_3_ for any rooted quartet tree can be estimated using the method-of-moments (MOM) estimators of Kubatko and Chifman (2020) given the value of *θ*. In order to estimate speciation times using the MOM estimators from the sequence data, the observed 15 site pattern frequencies of a quartet acquired from the data should be converted to observed site pattern probabilities that can be computed using (18) and (19) in Appendix C for 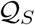 and 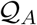, respectively.

While the MOM estimators are limited to *n* = 4, we can extend them to larger topologies by averaging *τ_y_* over all quartets that include node *y*. Let the three internal nodes in a quartet *q* be designated by *a_q_*, *b_q_*, and *c_q_* in the order of a postorder traversal. The mean age estimate for any node *y*, 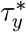, of the full tree over 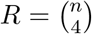 quartets is computed by:

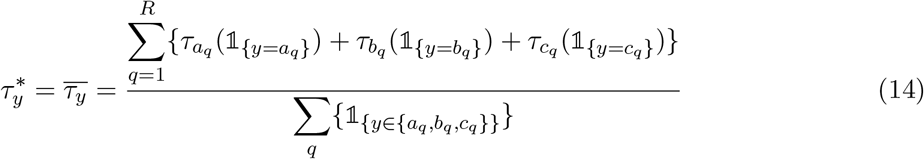

where 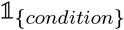 is an indicator function equal to 1 when *condition* is true and 0 otherwise, and summations are taken over all possible quartets. The denominator is just the number of quartets for which node *y* is included in the subtree induced by the *q*th quartet. In some cases, particularly when the taxa of interest have a complex evolutionary history, the estimate 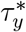 can be larger than the estimate 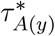. Biologically, this is impossible because it implies that the child existed before the parent. In this case, PhyNEST arbitrarily sets 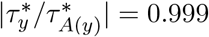 to use it as the starting point for optimization. The same procedure is applied when 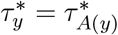.

In order to get good starting values for *τ* and *θ*, we first obtain a starting value for *θ* by minimizing the one-dimensional composite likelihood function corresponding to the negative logarithm of (11) viewed as a function of *θ* alone:

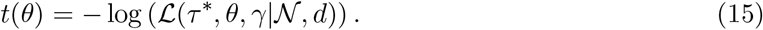

The set of node ages *τ*^*^ is computed using the MOM estimator described in (14) for any given value of *θ* with fixed *γ* (i.e., the user-specified value in the input network). In case no *γ* is provided, *γ* = 0.5 is arbitrarily selected. Minimization of *t*(*θ*) is performed using a one-dimensional, derivative-free method, with the initial *θ* value chosen as

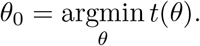

We begin with an *ad hoc* method that finds a loose interval (*A, B*) that contains *θ*_0_ using the idea that MOM estimators for realistic values of *θ*_0_ should be non-negative for all quartets in 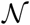. In this case, we say *θ*_0_ is *feasible*. First, we choose a reasonable lower bound value that is assumed to be less than *θ*_0_ (currently set as *A* = 10^−5^ in PhyNEST). Then the upper bound *B* is iteratively increased by a factor of *z*, where *z* ∈ (1, ∞) until *θ* is no longer feasible. We select the largest feasible *B* value evaluated, yielding an initial interval (*A, B*) with *A* ≤ *θ*_0_ ≤ *B*. Currently, *z* = 2 as default in PhyNEST.

Once this loose bound on the feasible region of *θ* is computed, we initiate a golden section search (Gill et al., 1981) to further shrink the interval by progressively narrowing the range of values of the specified interval that locates the minimum. In brief, we iteratively narrow down the interval (*A, B*) determined in the *ad hoc* method above by the golden ratio 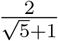 multiplied by the difference between the *A* and *B* values at that moment until |*B* − *A*| is smaller than a user specified tolerance value. At every iteration, we compute *t*(*A*) and *t*(*B*) (i.e., (15)) and keep a pair of values, either (*A, t*(*A*)) or (*B, t*(*B*)), whichever has smaller *t*(*θ*). If the selected pair happens to be (*A, t*(*A*)) we narrow the interval in the next iteration by reducing *B*; otherwise we narrow the interval by increasing *A*. One disadvantage of golden section search is that it converges relatively slow but PhyNEST overcomes this issue by setting a reasonably large tolerance value of 0.01 as default.

Finally, the minimization of (15) by optimizing *θ* is completed using the *localmin* procedure of Brent (2002) with the (*A, B*) bound computed from the golden section search algorithm. The determined *θ*_0_ is then used to set the starting node ages *τ*_0_ to the values of 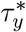 obtained using the method of (14), which are subsequently used as the starting values along with *γ* for the optimization of the composite likelihood function in (11).

### 3.3 Heuristic Search

#### 3.3.1 The moves

We search the space of phylogenetic networks to find a network that maximizes the composite likelihood. Instead of deriving new theoretical results on properties of network space, we implemented the following five moves (one operation to traverse the tree space and four operations for network space) using the relevant functions available in the Julia package PhyloNetworks (SolísLemus et al., 2017). Similar operations are used in the heuristics implemented in PhyloNet (Yu et al., 2014; Yu and Nakhleh, 2015) and in the SubNet Prune and Regraft (Bordewich et al., 2017) edit operation implemented in RF-NET 2 (Markin et al., 2022). Note that the input and output networks for these moves are semi-directed networks, hence these networks lack directionality except for the reticulation edges. The five moves are:

i. Nearest-neighbor interchange (NNI): Performed on a tree edge chosen uniformly at random that exchanges connectivity of the subtrees on the opposite sides of the chosen edge, within the main tree. Conducted when the current network is a tree (i.e., *h* = 0).
ii. Head move of a reticulation edge: Let *e* = (*u, v*) be a reticulation edge (i.e., *v* ∈ *V_N_*) of a network 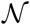 and let *b* be another edge in 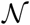. Let the vertex *v* be chosen at random from *V_N_* and denote the two incoming edges by *e*′ and *e*″. Select either *e*′ or *e*″ at random according to their inheritance probabilities (i.e., *e*′ has probability *γ* of being chosen and *e*″ has probability (1 − *γ*); we call the selected reticulation edge *e*). Define a new vertex *v*′ along a randomly-chosen edge *b* (i.e., *v*′ has in-degree one and out-degree one). The head of *e* is moved by replacing *v* with *v*′ so that *v*′ now has in-degree two and out-degree one (i.e., *v*′ ∈ *V_N_*). Node *v* is removed and the incoming and outgoing edges to *v* are merged.
iii. Tail move of a reticulation edge: Let *e* = (*u, v*) be a reticulation edge (i.e., *v* ∈ *V_N_*) of 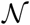 and let *b* be another edge in 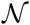. The vertex *v* is chosen at random and one of two incoming edges (*e*′ and *e*″) associated with *v* is selected according to their inheritance probabilities (i.e., *e*′ has the probability of *γ* to be chosen and *e*″ has (1 − *γ*); we call the selected reticulation edge as *e*). Define a new vertex *u*′ along a randomly-chosen edge *b* (i.e., *u*′ has in-degree one and out-degree one). The tail of *e* is moved by replacing *u* by *u*′ so that *u*′ has in-degree of one and out-degree two (i.e., *u*′ ∈ *V_I_*). Node *u* is removed and the incoming and outgoing edges to *u* are merged.
iv. Change the direction of a reticulation edge: Let *e* = (*u, v*) be a reticulation edge (i.e., *v* ∈ *V_N_*) of 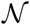. A vertex *v* is chosen at random from *V_N_* and one of the two incoming edges (*e*′ and *e*″) associated with *v* is selected according to their inheritance probabilities (i.e., *e*′ has probability *γ* of being chosen and *e*″ has probability (1 − *γ*); we call the selected reticulation edge *e*). The direction of *e* is ‘flipped’ by switching *u* and *v* thus *e* = (*v, u*), and *u* ∈ *V_N_* and *v* ∈ *V_I_*.
v. Insertion of a reticulation edge: When the number of reticulations is less than *h_max_* (the user specified maximum number of reticulations in the final network), two tree edges, *b*_1_ and *b*_2_, in 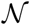 are selected at random. Two vertices with in-degree one and out-degree one, *u* and *v*, are created on *b*_1_ and *b*_2_, respectively. A reticulation edge *e* = (*u, v*) with length equal to zero and *γ* drawn uniformly from [0, 0.5) is created.

Any proposed network is checked to verify that it is of level 1, with *h* ≤ *h_max_*, and with at least one valid placement for the root. Since the computation of composite likelihood using (11) requires rooted quartets, we root every proposed network with a user-specified outgroup. In case there is no valid placement for the root using the chosen outgroup in the proposed network, the root is placed at random.

#### 3.3.2 Hill-climbing

Hill-climbing algorithm implemented in PhyNEST begins with an initial network (or tree) specified by the user. A robust and reliable starting topology (that is likely to be close to the true network) is expected to increase the chance of finding the global optimum (i.e., the topology with the highest composite likelihood in the given network space) with fewer steps. In case the confidence in the starting topology is low, PhyNEST has an option to conduct an NNI procedure on the specified starting topology at the onset of every run to initiate the search at the various (possibly better) points in the network space.

Denote the starting topology as 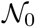 and let 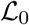 represent the negative logarithm of composite likelihood of 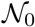. The procedure iteratively attempts to modify 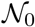 according to one of the possible five moves described above. We denote the modified network as 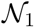 and its negative logarithm of composite likelihood as 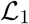. At each iteration, if 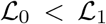, then 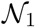 is discarded and a new modification is made to 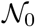 to create a new 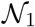 (i.e., failed movement); whereas if 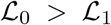, then 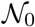 is replaced by 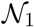 and a modification is made to the new 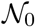 to create a new 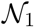. The algorithm will always accept a proposed network with a smaller negative logarithm of composite likelihood. The search terminates when either the number of iterations or the number of consecutive failed movements reaches a user-specified limit (the default is 10^5^ for iterations and 100 for failed movements in PhyNEST).

#### 3.3.3 Simulated annealing

We also implemented a simulated annealing algorithm (Černý, 1985; Aarts and Korst, 1989) in PhyNEST, as has been successfully applied in other phylogenetic settings (e.g., Barker, 2004; Stamatakis, 2005; Strobl and Barker, 2016; Salter and Pearl, 2001) for searching for the global optimum, as hill-climbing is only guaranteed to find local optima. Note many of the choices made in this section are motivated by Salter and Pearl (2001). Similar to the hill-climbing algorithm, the network space search is initiated from the user specified 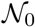 that has the negative logarithm of composite likelihood 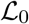. The algorithm makes the first step by modifying 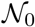 to create 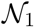 through one of the five moves described above, with the negative logarithm of composite likelihood 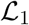. At each iteration, if 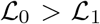, the algorithm always accepts 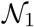 and replaces 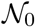 by 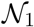. If 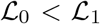, however, the algorithm accepts 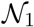 with a probability *t* given by:

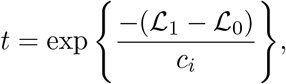

where *c_i_* is a control parameter described below. The probability of acceptance of an 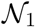 with a larger 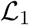 depends on how much worse 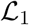 is compared to 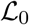 and decreases as the algorithm proceeds. In particular, this probability is controlled by the cooling schedule (Lundy, 1985; Lundy and Mees, 1986) with control parameter *c_i_* that is updated at each iteration *i* by setting:

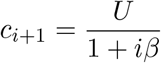

where *U* is an upper bound on the change in the composite likelihood in one iteration and *β* is the parameter that controls the rate of cooling. We estimate a reasonable *U* by running the algorithm for an initial ‘burn-in’ period. Specifically, a single move from 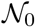 is made to create 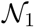, and 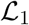 is computed, and the difference between 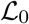 and 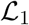 is recorded. Then we set 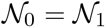 and repeat this procedure a user-specified number of times (25 times is the default in PhyNEST). Once this period has been completed, we set *U* to the maximum change in the negative log composite likelihood observed, and we set *β* to be:

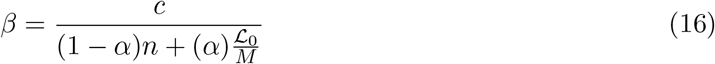

where *n* is the number of taxa, *M* is the length of the alignment, *c* and *α* are constants between 0 and 1, and 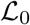 is the negative log composite likelihood of the starting topology. The motivation for setting *β* this manner is that the difficulty of the problem is influenced by both the number of taxa in the network and the magnitude of the negative log likelihood attributed to each site. The interval for *β* is [0, 1], where *β* ≈ 1 will result in faster cooling but an increased chance of getting trapped in local maxima, whereas *β* ≪ 1 will result in slower cooling but longer run times. The value of *β* is related to the reciprocal of a linear combination of *c* and *α*, where the constant *c* in the numerator allows the user to alter the rate of cooling without changing the particular linear combination selected in the denominator. PhyNEST sets *c* = 0.8 and *α* = 0.9 by default based on simulation study (see Results), although cautious selection of these constants by the user is strongly recommended.

The search terminates when the number of steps reaches the user-specified limit (currently set at 10^5^ iterations) or the number of iterations since the proposal of an uninvestigated topology reaches the user-specified upper bound. This upper bound is directly derived from the work of Salter and Pearl (2001) who converted the probability that an internal node would not have been selected during the set of proposed movements to a bound on the number of iterations under an NNI-move strategy for trees. We set the probability to a very small value, such as 9.5 × 10^−45^, by default. Note that the hill-climbing algorithm returns a point estimate of the network with the highest composite likelihood found by the algorithm. For simulated annealing, the algorithm constructs a list of the *k* trees of highest composite likelihood found during the search, where the number of trees included in the list is specified by the user (the default is *k* = 10).

## 4 Results

### 4.1 Simulations

#### 4.1.1 Finding optimal *α* and *c* for the simulated annealing algorithm

The cooling schedule is a crucial component of the simulated annealing algorithm and the initial values of the control parameters *α* and *c* in (16) must be specified prior to the network search. While Salter and Pearl (2001) reported *α* = *c* = 0.5 seemed to work well when searching the space of phylogenetic trees, the effect of these parameters when searching the space of phylogenetic networks is unknown. Using simulation under a simple scenario, we explore the effect of *α* and *c* in terms of efficiency and accuracy when searching the network space.

We selected a phylogenetic network with *n* = 5 and *h* = 1 (Fig. 2a) as a model to generate data. Let the speciation times *τ_j_* be indexed in a postorder traversal by *j* = 1, 2, …, 5. The speciation times were set in coalescent units (CUs) to *τ*_1_ = *τ*_2_ = 1.5, *τ*_3_ = 3, *τ*_4_ = 5, and *τ*_5_ = 10. We set *γ* = 0.4 for the reticulation edge between species 1 and 2 and *θ* = 0.02. We generated 10^5^ gene trees using *ms* (Hudson, 2002) (i.e., 0.4×10^5^ for one of the parental trees that forms a cherry with species 1 and 2 and 0.6 × 10^5^ for the other) followed by the simulation of DNA sequences using *seq-gen* (Rambaut and Grass, 1997) under the JC69 model with a length of 100 base pairs (b.p.) per locus to mimic the short-read lengths in next-generation sequence methods (Kubatko and Chifman, 2019). All sequences were formatted using *goalign* (https://github.com/fredericlemoine/goalign) and *snp-sites* (Page et al., 2016).

**Figure 2:**
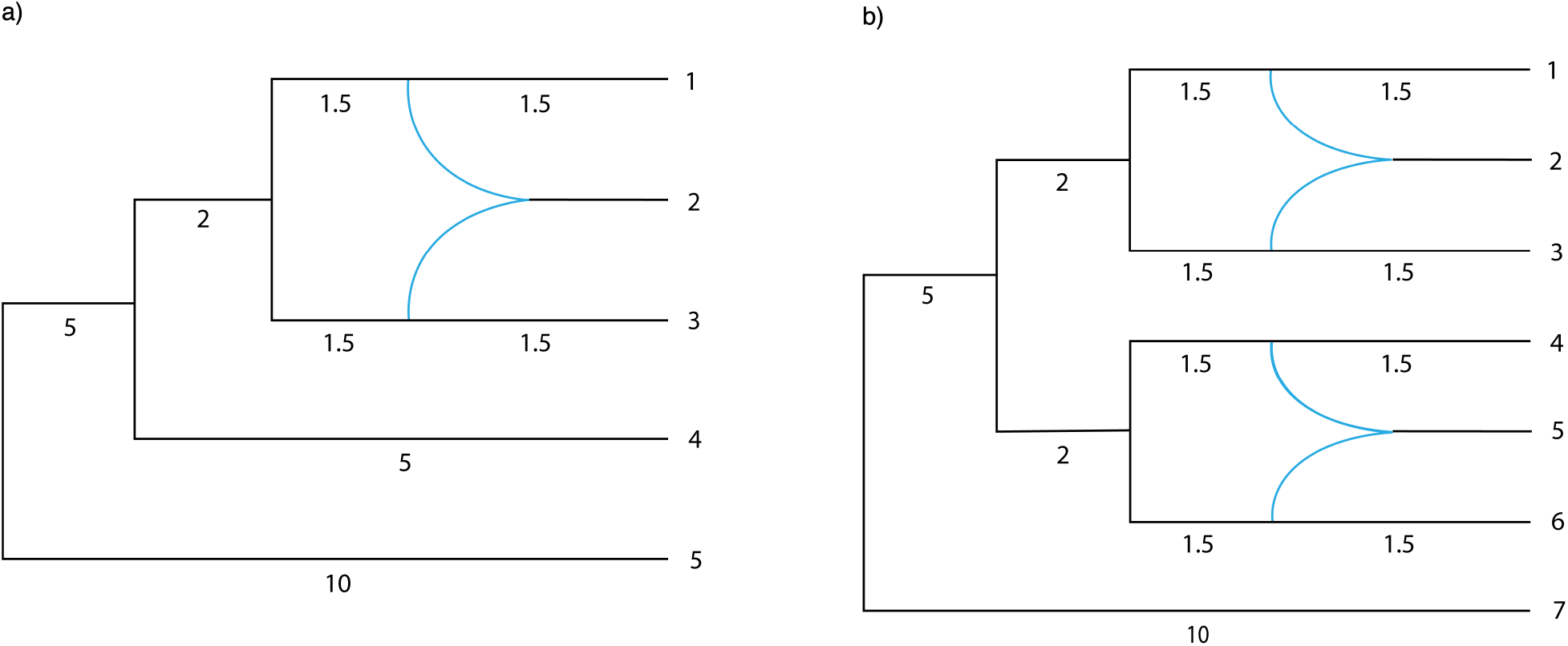
Hybridization scenarios with (a) five tips and a single reticulation and (b) seven tips and two reticulations. Node ages are in coalescent units.

We selected *α, c* ∈ {0.01, 0.1, 0.2, 0.3, 0.4, 0.5, 0.6, 0.7, 0.8, 0.9, 0.99}, producing 121 combinations for evaluation. For each pair of *α* and *c*, 20 independent runs were executed using the simulated annealing algorithm in PhyNEST. We initiated the search using a random starting topology with the correctly specified outgroup taxon and the correct number of maximum reticulations (i.e., *h_max_*=1). The maximum number of steps for each run was set to 10^3^ to prevent an unnecessarily prolonged search. At the completion of each independent run, we recorded the number of steps taken and determined if the true network was reconstructed. For each pair of *α* and *c*, we computed the efficiency-accuracy score (*F*) using:

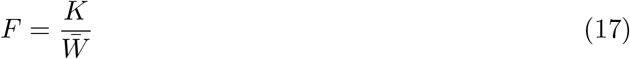

where 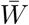 is the average number of steps and *K* is the number of times the selected final network is isomorphic to the true network for the 20 runs. We interpreted higher values of *F* to indicate greater efficiency as the true network is found with fewer steps.

Figure 3a summarizes the number of steps at the termination of each run for every pair of *α* and *c*. Generally, the number of steps is inversely proportional to both *α* and *c* (i.e., more steps are required for smaller *α* and *c*). The search almost always required the maximum number of steps when *c* = 0.01 for *α* < 0.99, *c* = 0.1 when *α* < 0.7 and *c* = 0.2 when *α* < 0.2. Note that in most of these cases, the search ended because the maximum number of steps was reached, not because it converged to an optimum, thus a higher threshold may be necessary when using smaller values of *α* and *c*. The search never required 10^3^ steps when *α* = 0.99 regardless of *c* and the search required ≈ 78 steps on average when *α* = *c* = 0.99. The search required fewer than 500 steps when *α* < 0.2 with *c* < 0.3.

**Figure 3:**
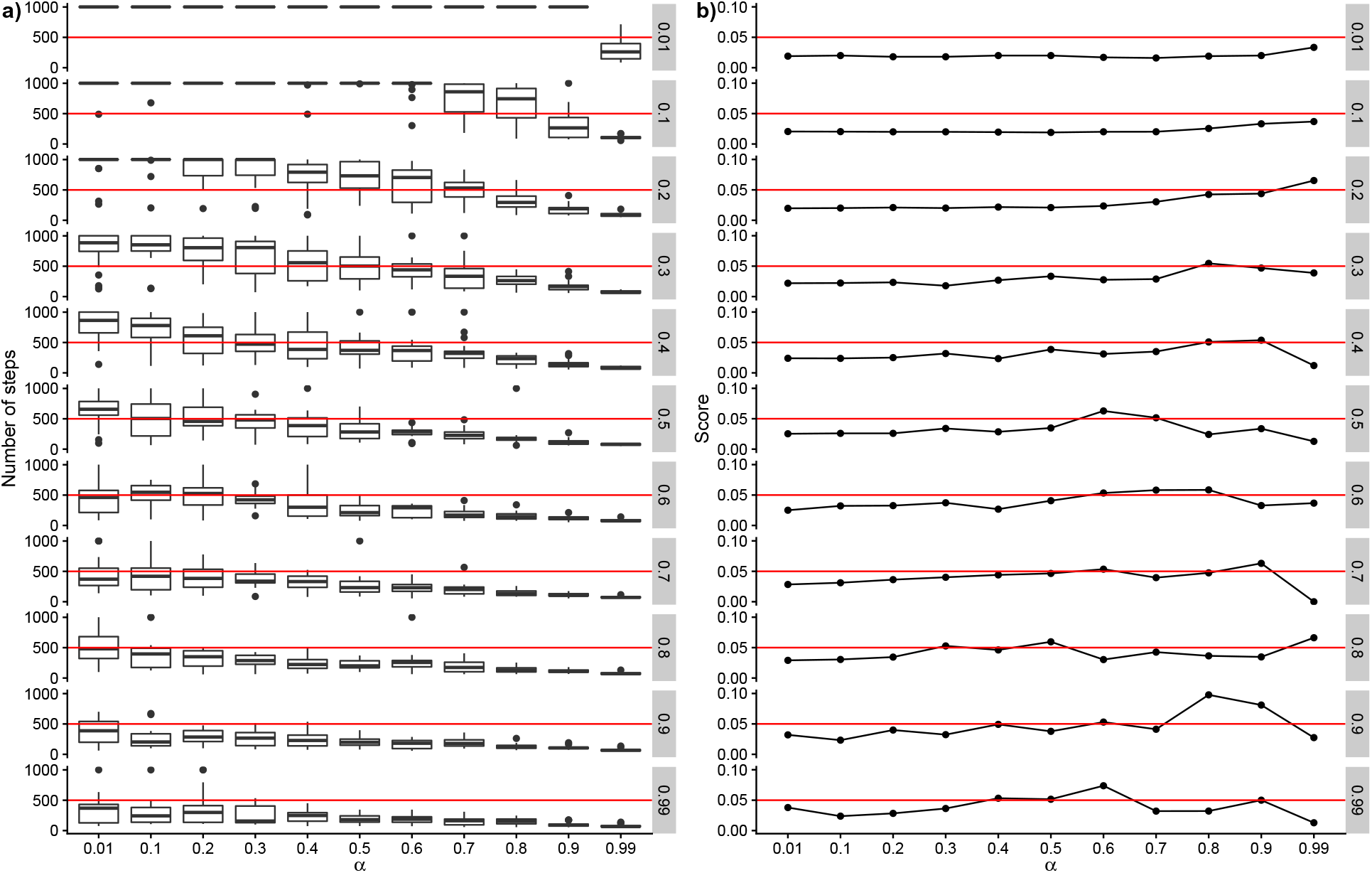
(a) Each box plot summarizes the number of steps taken at the point of termination of the 20 independent runs for each combination of *α* and *c* during network search using the simulated annealing algorithm. We selected *α, c* 0.01, 0.1, 0.2, 0.3, 0.4, 0.5, 0.6, 0.7, 0.8, 0.9, 0.99, hence 121 combinations were evaluated here. The left y-axis represents the number of steps, and ranges from 0 to 1000 (threshold), the x-axis represents tested values of *α*, and the value in the gray boxes on the right y-axis represents *c*. The red solid horizontal line in each pane indicates 500 steps. (b) Line graphs showing the computed *F* score using Equation (17) for the 121 combinations of *α* and *c* evaluated. The y-axis on the left represents the scores, the x-axis represents tested values of *α*, and the values in the gray boxes on the y-axis on the right represent tested values of *c*. The horizontal red line for each pane indicates the score of 0.05. The highest score was observed when (*α, c*)=(0.8,0.9) followed by (0.9,0.9). We selected (*α, c*)=(0.8,0.9) as the default values for PhyNEST when using the simulated annealing algorithm.

Figure 3b summarizes *F* scores using (17) for each pair of *α* and *c*. The scores were arguably low when *α* ≤ 0.7, but were the highest when *c* = 0.9 with *α* = 0.8 and 0.9. Using this result, we set the default values of *α* and *c* as 0.8 and 0.9, respectively, to initiate the network search using the simulated annealing algorithm in PhyNEST. However, it is difficult to make a concrete generalization based on the evaluations taken here because of the stochastic nature of the network searching process. Also, our test is based only on a very simple scenario and extensive evaluation using various scenarios is needed. In practice, it is strongly recommend that users set *α* and *c* to relatively low values (e.g., *α* = *c* = 0.25) with a large maximum number of steps depending on the dataset size and the availability of computational resources to increase their chance of finding the global optimum.

#### 4.1.2 Efficiency of parsing the input sequence alignment

This section provides an approximate idea on the running time for parsing the input sequence alignment using various combinations of number of sequences *n* and the alignment length *M*. ‘Parsing’ includes reading in the sequence alignment, computing the site pattern frequencies for 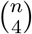 quartets, and rearranging the computed site patterns for 4! permutations for each quartet decomposed from 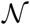 (see Table 3 in Appendix B). Following the data simulation procedure described in the previous section, we generated datasets that have *n* ∈ {4, 5, …, 15} with *M* from 5 × 10^3^ to 10^6^ b.p. in increments of 5 × 10^3^, resulting in 240 combinations of *n* and *M*. Since the relationships among taxa are not of interest in this section, we used two very simple scenarios. Scenario 1 contains two species, written as (A,B); in Newick format, with *τ*_1_ = 2, where two and *n* − 2 individuals belong to Species A and B, respectively. Scenario 2 contains four species, written (A,(B,(C,D))); in Newick format, with *τ*_1_ = 4, *τ*_2_ = 10, and *τ*_3_ = 20 indexed in a postorder traversal, where each species contains a single individual except for species B which contains *n* − 3 individuals. We recorded the running time taken for each dataset on a personal laptop with 7.62 gigabyte of RAM and 2 processors.

We plotted the running time for parsing the alignments generated under Scenarios 1 and 2 in Figures 4a and 4b, respectively. For each *n*, we drew a smooth curve to enhance visibility using local polynomial regression. The running time taken to parse the data increased exponentially with dataset size in both scenarios. In Scenario 1, the shortest running time (≈ 0.15 seconds) was observed when *n* = 4 and *M* = 5 × 10^3^, but even when *n* and *M* were increased to 15 and 10^6^, respectively, it took ≈ 30 seconds. In Scenario 2, running time was the smallest when *n* = 4 and *M* = 5 × 10^3^ (≈ 0.42 seconds) and the greatest when *n* = 15 and *M* = 10^6^ (≈ 432.43 seconds). We noticed the time required to parse the data was greater in Scenario 2 than in Scenario 1, given the same *n* and *M*. This is because the sequences are more divergent from each other in Scenario 2, and a divergent dataset results in larger matrices and vectors when summarizing the sequences, requiring longer computation times. Nevertheless, this is orders of magnitude faster than summarizing the alignment into a series of gene trees for use as input to summary-based network inference methods.

**Figure 4:**
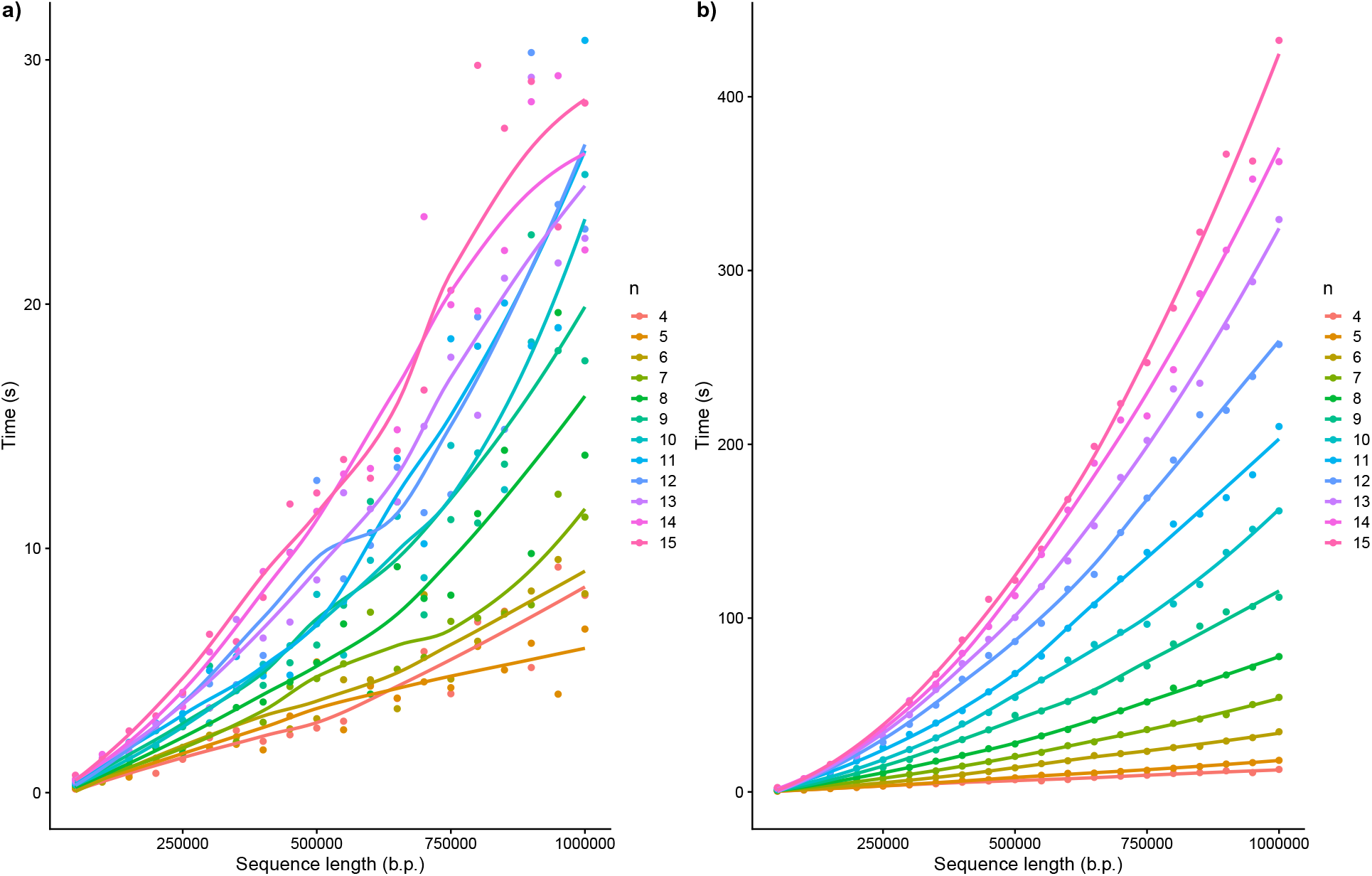
The impact of dataset size on the computational time taken to parse the input dataset for (a) Scenario 1 and (b) Scenario 2. the y-axis represents time measured in seconds and the x-axis represents the sequence lengths in the number of base pairs. The number of individuals per dataset, *n*, is color coded as in the legend on the right side. A smooth curve was drawn for each *n* to enhance visualization.

#### 4.1.3. Accuracy of parameter estimation

This section presents results concerning the accuracy of parameter estimates of a network, *τ, γ*, and *θ*, that are optimized by maximizing the composite likelihood function (11). We considered the two hybridization scenarios in Figure 2. Let the speciation times *τ_j_* in each scenario be indexed in a postorder traversal *j* = 1, 2,…, *n* + *h* − 1. The speciation times in Scenario 1 were set in CU as *τ*_1_ = *τ*_2_ = 1.5, *τ*_3_ = 3, *τ*_4_ = 5, and *τ*_5_ = 10. Similarly for Scenario 2, the speciation times were set as *τ*_1_ = *τ*_2_ = *τ*_3_ = *τ*_4_ = 1.5, *τ*_5_ = *τ*_6_ = 3, *τ*_7_ = 5, and *τ*_8_ = 10. We set *γ* = 0.5 for every reticulation and *θ* = 0.02. For both scenarios, we simulated *g* gene trees where *g* ∈ {500, 1000, 5000, 10000} using *ms*. We additionally simulated *g* = 20000 for Scenario 2 since more data may be required for accurate parameter estimation for a larger and more complex scenario. We decomposed Scenario 1 and 2 into an array of parental trees and simulated 0.5 × *g* and 0.25 × *g* gene trees for each parental tree in Scenario 1 and 2, respectively. The simulated gene trees were then used as input to simulate DNA sequences using *seq*-*gen* with a length of 100 b.p. per locus under the JC69 model. We used 100 replicates for each *g*.

Figure 5 presents estimates of *τ, γ*, and *θ*, for various *g* in Scenario 1 (Fig. 5a–c, respectively) and Scenario 2 (Fig. 5d–f, respectively). Note that in Figures 5a and 5d, we computed Δ*T*, the difference between true and the estimated *τ*, to enhance readability. For all estimates in both scenarios, differences between the true and estimates and the variances between the replicates reduced as the dataset size increase. The times of ancestral vertices (i.e., *τ*_5_ and *τ*_8_ in Scenarios 1 and 2, respectively) were more difficult to estimate than the times of recent vertices as reflected in the increased variance among the replicates and greater deviations from the true value. The estimates of *γ* and *θ* were exceptionally accurate in all *g*. We omitted two replicates for Scenario 2 when *g* = 500 as their estimates seemed infeasible (Δ*T* > 2000 CU and *θ* ≈ 0.0. Even in these cases, however, the estimates of *γ* were reasonably accurate with (*γ*_1_, *γ*_2_) = (0.56, 0.61) for one omitted replicate and (0.59, 0.62) for the other.

**Figure 5:**
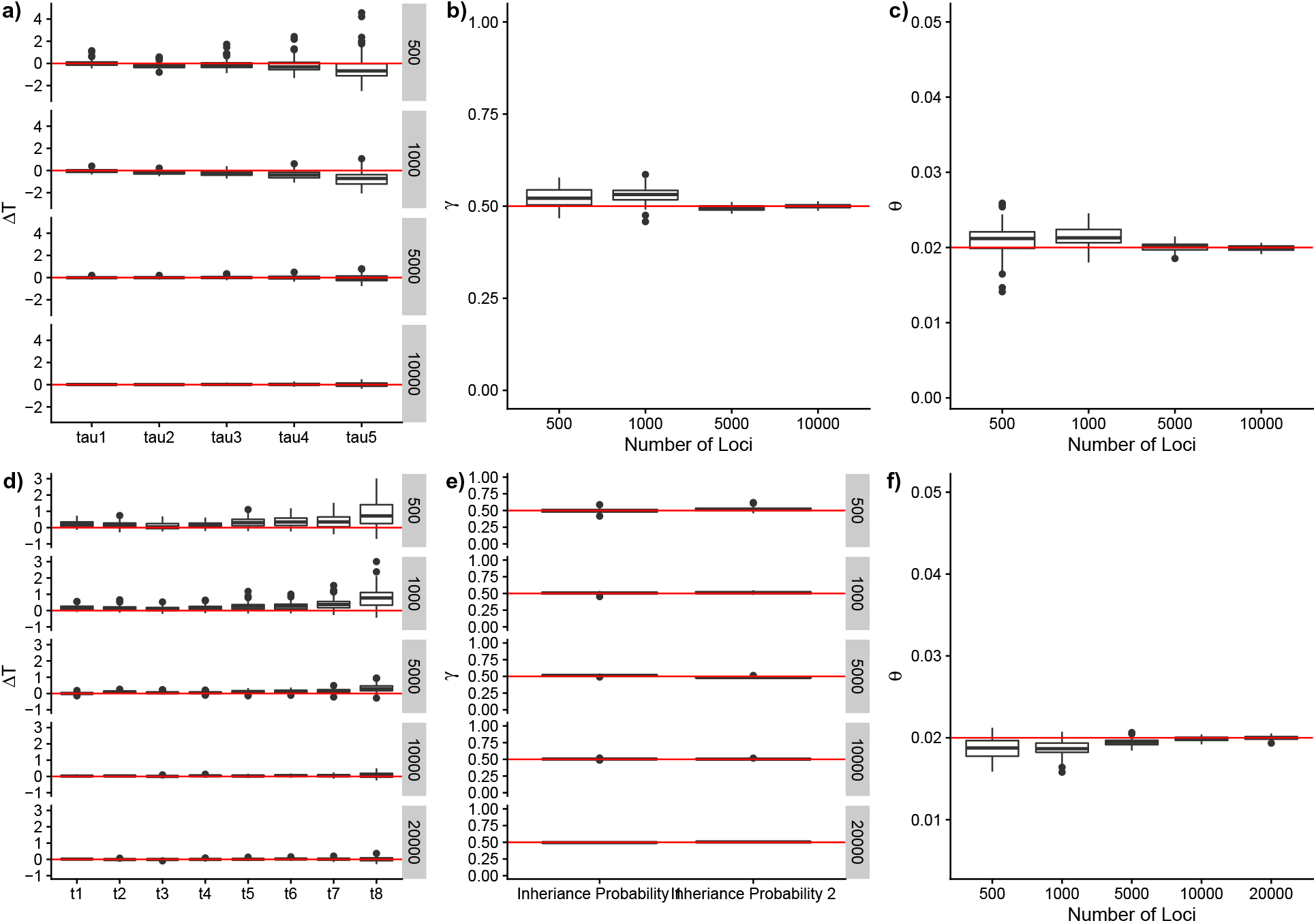
(a) Boxplots that show the difference between the estimates of the speciation times (*τ*) in CU and true *τ* (Δ *T*; y-axis on the left) for different sized datasets for Scenario 1 and (d) Scenario 2. The y-axis on the right shows the number of gene trees simulated. The red horizontal line indicates Δ*T* = 0 (i.e., the estimate and the true value are identical). (b) Boxplots that show the estimates of the inheritance probability (*γ*; y-axis) that optimize the composite likelihood function for Scenario 1 and (e) Scenario 2, for different sized datasets (x-axis in (b) and gray boxes on right y-axis in (e)). On the x-axis of (e), *γ*_1_ represents the probability associated with hybrid taxon 2 and *γ*_2_ represents for hybrid taxon 5. The red horizontal line indicates *γ* = 0.5, the true value of *γ*. (c) Boxplots that show the estimates of the effective population size parameter (*θ*; y-axis) that optimizes the composite likelihood function for Scenario 1 and (f) Scenario 2, for different sized datasets. The red horizontal line indicates *θ* = 0.02, the true value of *θ*.

#### 4.1.4 Comparative performance of PhyNEST with other methods

In this section, we used simulation to examine the question of whether PhyNEST can return networks that are competitive with respect to topological accuracy and computational efficiency with two existing methods, SNaQ implemented in the Julia package PhyloNetworks and the MPL function implemented in PhyloNet (SNaQ and PhyloNet hereafter, respectively), that also use a composite likelihood framework. We simulated sequence alignments from two network topologies, 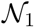 and 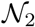 (Fig. 2a and 2b, respectively), where 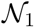 has *n* = 5 with *h* = 1 and 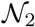 has *n* = 7 with *h* = 2, where *n* and *h* represent the number of tips and the number of reticulation(s) in the network, respectively. Let the speciation times *τ_j_* in each 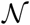 be indexed in a postorder traversal with *j* = 1, 2,…, *n* + *h* − 1. The speciation times in 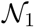 were set in CU. as *τ*_1_ = *τ*_2_ = 1.5, *τ*_3_ = 3.0, *τ*_4_ = 5.0, and *τ*_5_ = 10.0. Similarly, the speciation times in 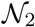 were set as *τ*_1_ = *τ*_2_ = *τ*_3_ = *τ*_4_ = 1.5, *τ*_5_ = *τ*_6_ = 3.0, *τ*_7_ = 5.0, and *τ*_8_ = 10.0. We set *γ* = 0.5 for every reticulation and *θ* = 0.003 for all scenarios. Let *g* be the number of loci. For 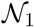, we simulated *g* = {50, 200, 500, 1000, 3000, 5000} gene trees using *ms*. Since *ms* only takes input in a tree format and cannot be used to simulate gene trees directly from a network, we decomposed 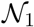 into an array of parental trees and simulated 0.5 × *g* gene trees for each parental tree. For 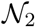, we simulated *g* = {40, 200, 500, 1000, 3000, 5000} gene trees using *ms*. We decomposed 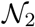 into an array of parental trees and simulated 0.25 × *g* gene trees for for each parental tree. For each gene tree, we generated a multiple sequence alignment of 250 b.p. using *seq-gen* under the JC69 model.

For each locus, we estimated an unrooted gene tree using IQ-TREE (Nguyen et al., 2015) with the DNA substitution model specified to JC69 and allowed automatic selection of the number of CPU cores for the computation. Using the set of estimated *g* gene trees from the entire sequence alignment, we computed a table of quartet concordance factors (CFs) using PhyloNetworks to use as input for network inference with SNaQ. Then, we rooted all gene trees using the true outgroup for use as input to PhyloNet. Finally, we used *goalign* and *snp-sites* to create a single alignment in sequential PHYLIP format for input to PhyNEST.

Using the prepared input files, we estimated the network using SNaQ, PhyloNet, and PhyNEST using both the hill-climbing and the simulated annealing heuristics (referred to as PhyNEST HC and PhyNEST SA, respectively hereafter), each with 10 independent runs. One of the estimated gene trees was randomly selected as a starting topology for SNaQ and PhyNEST analyses whereas for PhyloNet analysis, we allowed the search to be initiated using the optimal species tree under the minimizing the number of deep coalescences criterion (Than and Nakhleh, 2009) as suggested by the developers (Yu and Nakhleh, 2015). The starting topology for SNaQ and PhyNEST, however, was modified using NNI rearrangement in each independent run. We used the default settings for SNaQ and PhyloNet analyses. For PhyNEST SA we set *α* = *c* = 0.25. For all analyses, we specified the true number of reticulations as the expected number of reticulations in the estimated network. Accuracy of the estimated network obtained from each approach was assessed by measuring the distance between the true and estiamted networks using tree-, cluster-, and tripartition-based distance metrics, implemented in PhyloNet. For all three metrics, the estimated network is topologically closer to the true network when the distance metric is closer to zero. We also recorded the time taken for gene tree estimation using IQ-TREE and the running time for each analysis. We conducted 100 replicates for each *g*, although one or more analysis failed to complete in 1 of the 100 replicates when *g* = 1000 and 5000 and 2 of the 100 replicates when *g* = 3000 for 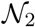 due to excessively long computation times. All analyses were carried out using the College of Arts and Sciences’ high-performance computing cluster (Unity) at The Ohio State University.

Figure 6a summarizes the running time for each replicate for the different values of *g* for 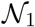. The computation times for all four methods tested were generally consistent across *g*. PhyloNet was the fastest, taking ≈ 25 seconds on average, followed by SNaQ with ≈ 45 seconds on average; PhyNEST HC required ≈ 7 minutes for computation, while PhyNEST SA took ≈ 4 hours. Figure 7a summarizes the time required for gene tree estimation for different values of *g* using IQ-TREE for 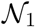. The estimation of a single gene tree took ≈ 94 seconds on average, which accumulated with increasing *g* and ranged from ≈ 77 minutes when *g* = 50 to ≈ 130 hours when *g* = 5000. Parsing the input alignments for PhyNEST required < 1 minute.

**Figure 6:**
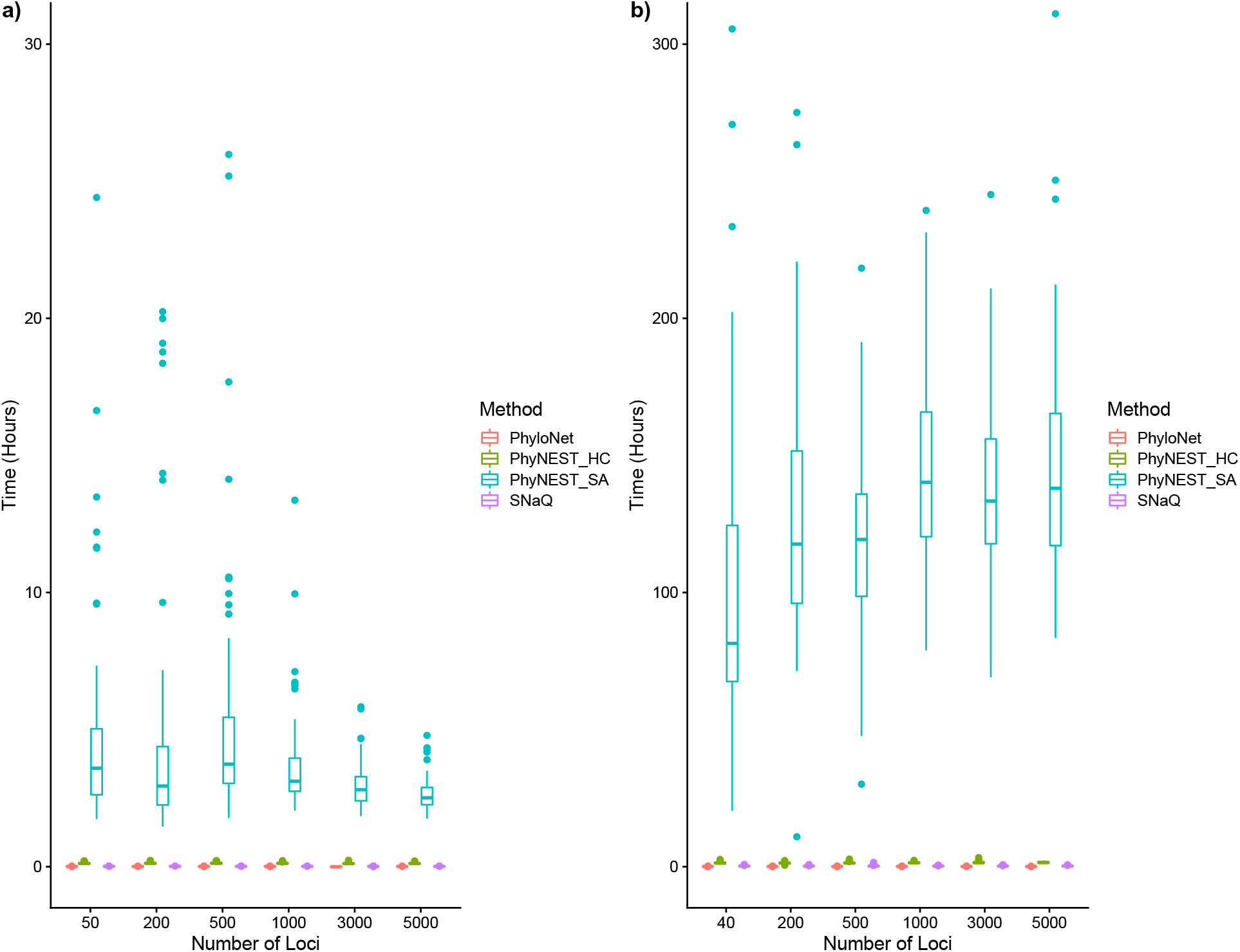
Summary of running time (y-axis) for the four methods evaluated, PhyloNet, SNaQ, PhyNEST using hill-climbing (PhyNEST HC), and PhyNEST using simulated annealing algorithm (PhyNEST SA), using simulated data for the hybridization scenario with (a) five leaves and one reticulation and (b) seven leaves and two reticulations over various numbers of loci (x-axis).

**Figure 7:**
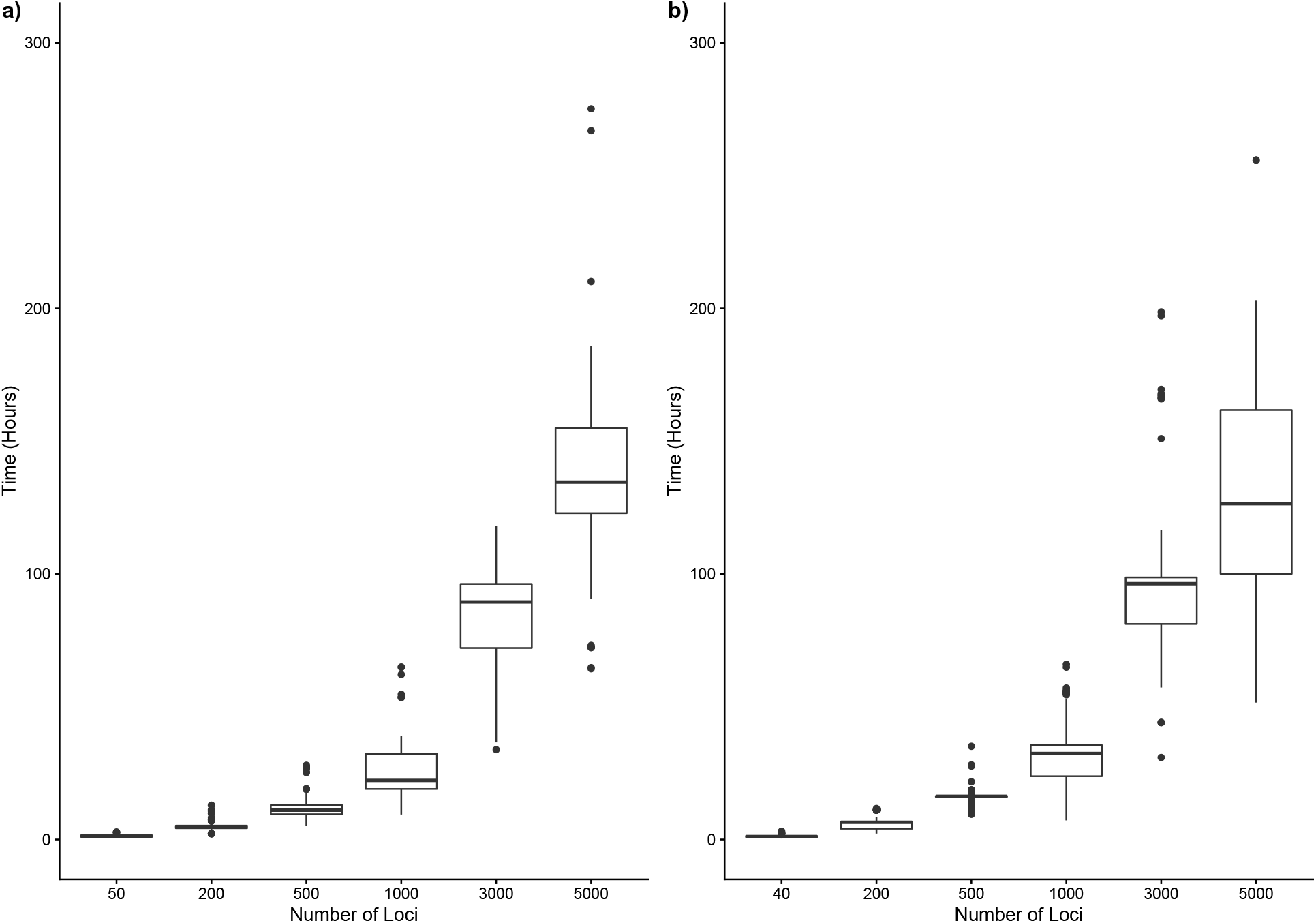
Summary of time taken for gene tree estimation (y-axis) using IQ-Tree over different number of loci (x-axis), using simulated data for the hybridization scenario with (a) five leaves and one reticulation and (b) seven leaves and two reticulations.

The tree- and cluster-based distance metrics (Fig. 8a, 8b) show that PhyNEST did an excellent job in reconstructing the true network when *g* > 500. Even when *g* < 500, the metrics for PhyNEST were closer to zero than for PhyloNet and SNaQ. It was not uncommon to observe PhyloNet and SNaQ estimating networks that were very distant from the true network when *g* = 50. Although this pattern disappeared when *g* < 500 for SNaQ, it persisted even when *g* = 3000 for PhyloNet. Based on the tripartition-based distance metric (Fig. 8c), when *g* = 50, PhyloNet and SNaQ rarely reconstructed the true network, whereas PhyNEST HC and PhyNEST SA recovered it much more frequently. Here, some of the networks estimated by SNaQ showed the greatest distance from the true network compared to the other three approaches. The accuracy of the estimates for the different methods were similarly good when *g* = 200, 500, and 1000, where the frequency of recovering the true network gradually increased with *g*. Note that when *g* = 1000, all methods estimated a network that is isomorphic to the true network in about half of the replicates. When *g* = 3000, all methods performed similarly, with SNaQ reconstructing the true network the most frequently, and when *g* = 5000 all methods almost always correctly identified the true relationships. Although assessment of each of the methods based on the three metrics generally followed similar patterns, we question the reliability of tree- and cluster-based distance metrics as they seem excessively liberal, yielding a distance of zero even when the two topologies being compared were not isomorphic. For example, tree- and cluster-based distances of zero were computed (tripartition-based distance=0.5) when the estimated network can be written as

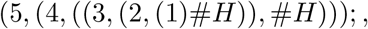

a scenario where species 1 is the hybrid of species 2 and the common ancestor of species 2 and 3, is compared with 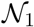. In this network scenario, the hybrid taxon is incorrectly identified and has different biological interpretations from 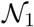. Understanding the cause of this issue is beyond the scope of this study, but we suspect this is possibly due to the fact that the set of parental trees for both network are the same.

**Figure 8:**
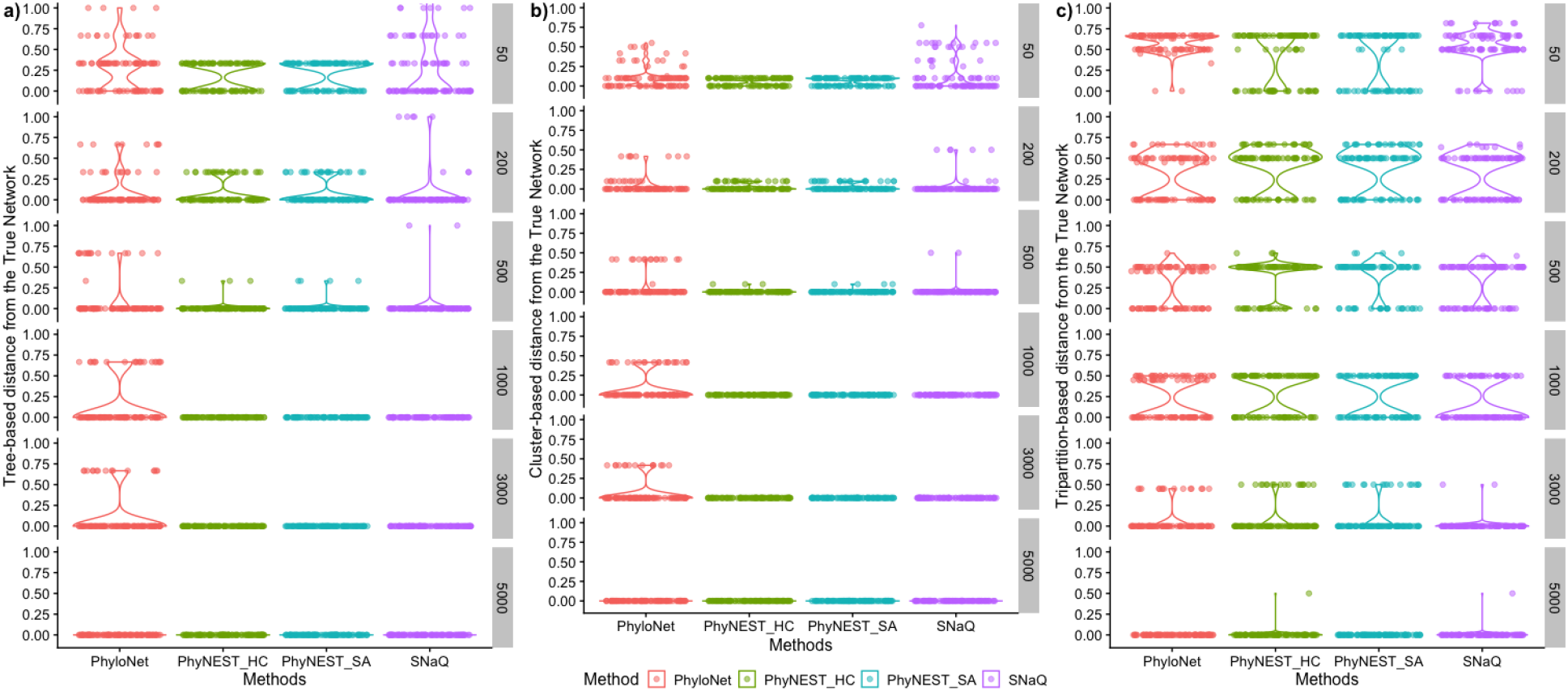
Comparison of methods with respect to network topological accuracy using simulated data for the hybridization scenario with five leaves and one reticulation. The accuracy was measured using the (a) tree-, (b) cluster-, and (c) tripartition-based distance metric over various numbers of loci (gray boxes on right). Each dot represents one of 100 replicates made for each locus size using one of four methods. The underlying violin plot shows the frequency of the computed distance.

Figure 6b summarizes the running time for each replicate for different numbers of genes evaluated for 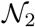. Similar to the case of 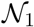, the computation times were consistent regardless of *g*, taking < 10 minutes for SNaQ and PhyloNet, and ≈ 100 minutes and ≈ 110 hours on average for PhyNEST HC and PhyNEST SA, respectively. Figure 7b summarizes the time required for gene tree estimation for different numbers of trees using IQ-TREE for 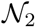. The estimation of a single gene tree took ≈ 120 seconds on average, which accumulated with increasing *g*, ranging from ≈ 61 minutes when *g* = 40 to ≈ 150 hours when *g* = 5000. Again, parsing the input alignments for PhyNEST was a matter of minutes.

Figures 9a and 9b present tree- and cluster-based distance between the estimated and true networks for each replicate for different numbers of genes for 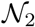. Both plots show that PhyNEST was more frequent in reconstructing the true network than both SNaQ and PhyloNet when *g* ≤ 500, and all methods seem to perform similarly when *g* ≥ 1000. Focusing on the tripartition-based distance metric in Figure 9c, all methods generally performed similarly well. The frequency of reconstructing the true network gradually increased with *g*. When *g* = 5000, PhyNEST SA recovered the true network in every replicate, showing the greatest accuracy among the four methods evaluated.

**Figure 9:**
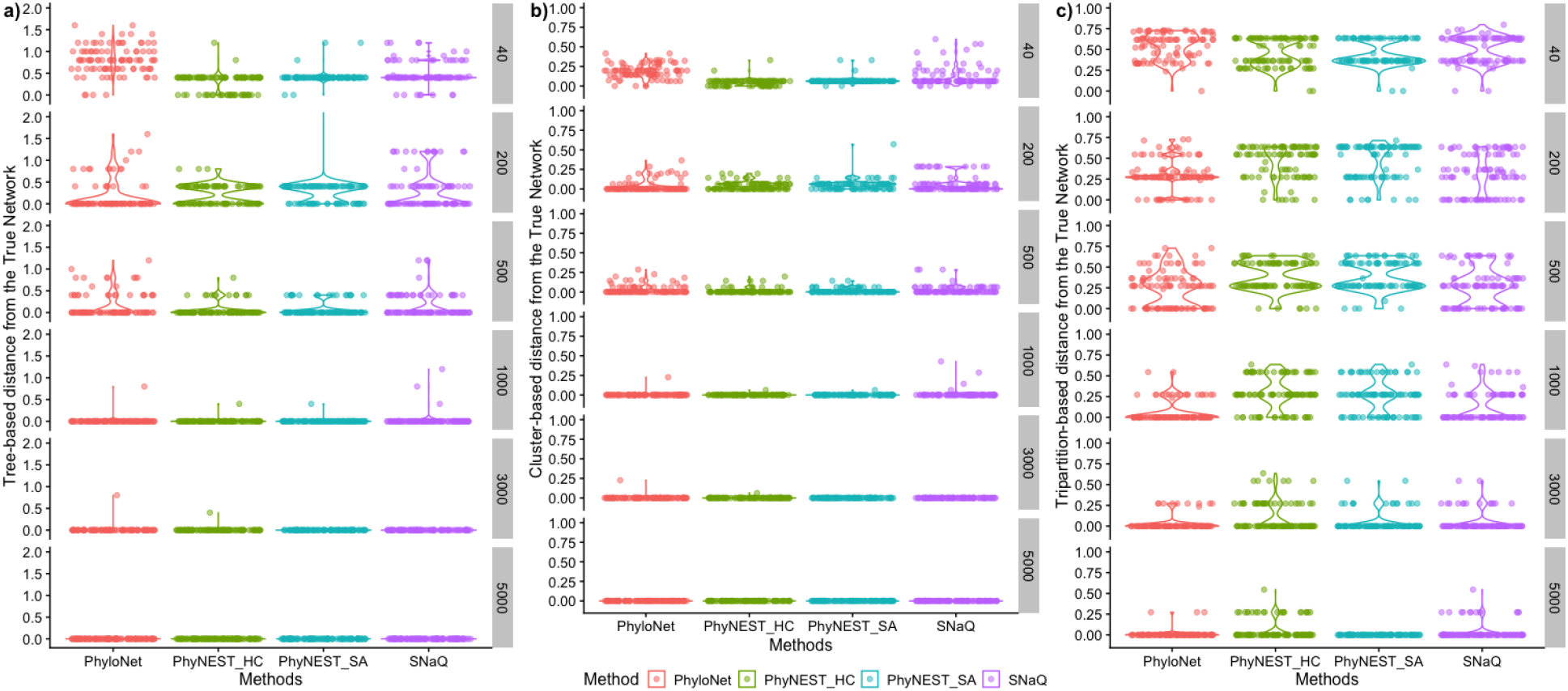
Comparison of methods with respect to network topological accuracy using simulated data for the hybridization scenario with seven leaves and two reticulations. The accuracy was measured using the (a) tree-, (b) cluster-, and (c) tripartition-based distance metric over various numbers of loci (gray boxes on right). Each dot represents one of 100 replicates made for each locus size using one of four methods. The underlying violin plot shows the frequency of the computed distance.

### 4.2 Empirical Data

#### 4.2.1 Hybrid speciation in *Heliconius* butterflies

We analyzed whole-genome sequences from Martin et al. (2013) to reconstruct the evolutionary history of four *Heliconius* butterflies. Using 31 individuals composed of six ingroup and four outgroup species, we created two sets of alignments, each of which contains four sequences from four species: (1) *Heliconius melpomene rosina* (ro2071), *H. m. melponeme* collected in French Guiana (meF13435), *H. cydno* (chi553), and *H. pardalinus* (Par371; outgroup) and (2) *H. timareta* (tiP86), *H. m. amaryllis* (am216), *H. m. melponeme* collected from French Guiana (meF13435), and *H. pardalinus* (Par371; outgroup). We note the sequence names in the original alignment in parenthesis. Each sequence in both alignments contained 248,822,400 nucleotides. The species in each dataset were selected based on Figure 2 in Martin et al. (2013) where strong gene tree–species tree discordance was observed based on the set of maximum likelihood gene trees estimated for non-overlapping 100-kilobase pair windows throughout the genome.

PhyNEST was initiated with the starting topologies specified for alignments (1) as:

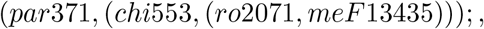

and (2) as:

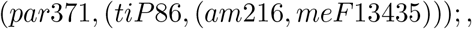

repsectively, based on the species tree topology provided in Martin et al. (2013). For each alignment, both hill climbing and simulated annealing were conducted for *h_max_* = 1. For simulated annealing, we set *α* = *c* = 0.25. For alignment (1), it took 212.779 seconds for the hill climbing search and 290.827 seconds for simulated annealing and for alignment (2), it took 144.716 seconds for the hill climbing search and 257.566 seconds for simulated annealing.

The estimated network using alignment (1) showed identical relationship to the analyses of Martin et al. (2013) with *H. m. rosina* being the hybrid of *H. cydno* and *H. m. melpomene* from French Guiana (Fig. 10a). In this dataset, PhyNEST estimated *γ* = 0.524, representing the proportion of the genome inherited from *H. m. melpomene* to *H. m. rosina*. This value was very close to the proportion of gene trees in the genome that matched the species tree topology (=53.0%). PhyNEST estimated speciation times *τ_j_*, indexed by a postorder traversal, *j* = 1, 2, 3, 4, as *τ*_1_ = 4.08 × 10^−19^, *τ*_2_ = 1.34 × 10^−18^, *τ*_3_ = 0.13135, and *τ*_4_ = 0.27946. Very short estimated values for *τ*_1_ and *τ*_2_ support the interpretation in Martin et al. (2013) for recent interspecific gene flow. Similarly, the estimated network using alignment (2) aligned with the interpretations of Martin et al. (2013) with *H. m. amaryllis* being the hybrid of *H. timareta* and *H. m. melpomene* from French Guiana (Fig. 10b). Here, PhyNEST estimated *γ* = 0.344 for the reticulation edge from the ancestor of *H. m. melpomene* to *H. m. amaryllis* with speciation times *τ*_1_ = 3.00 × 10^−19^, *τ*_2_ = 3.80 × 10^−15^, *τ*_3_ = 0.33108, and *τ*_4_ = 0.54605.

**Figure 10:**
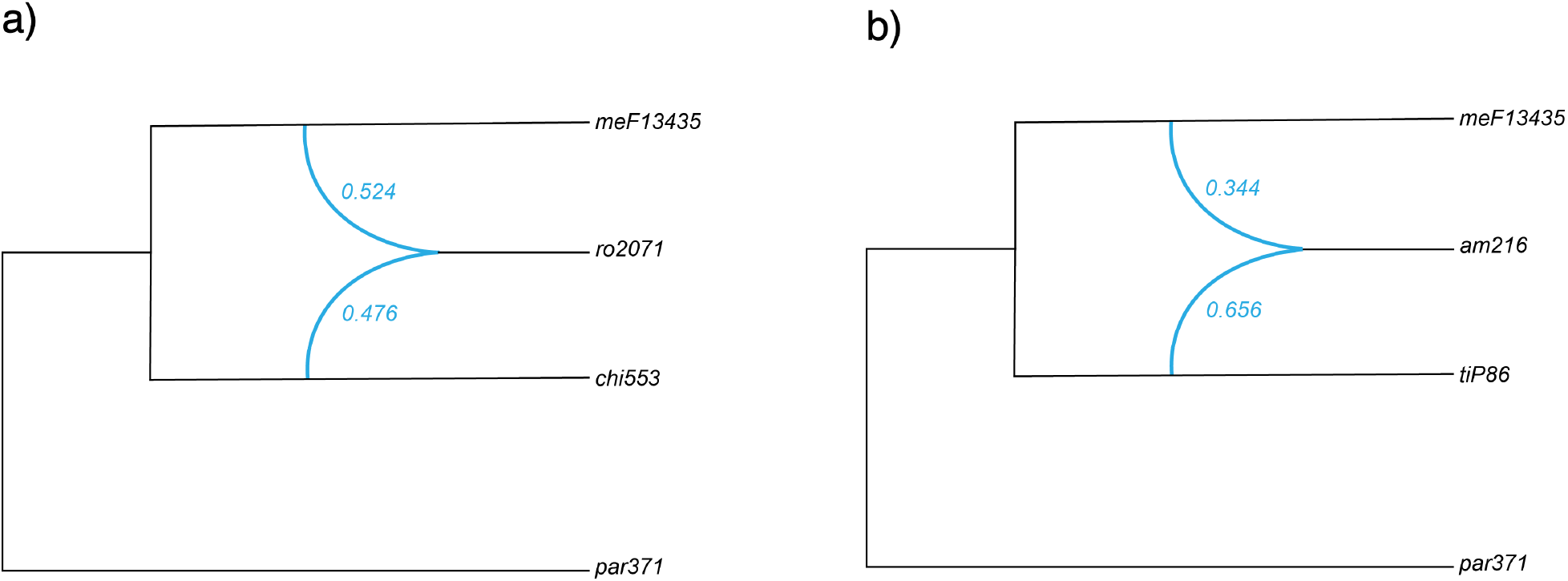
Hybrid speciation of *Heliconus* spp. inferred with PhyNEST from the sequence alignment of ≈ 250 million base pairs published in Martin et al. (2013). Each network contains four species, *Heliconius melpomene rosina*, *H. m. melponeme* collected in French Guiana, *H. cydno*, and *H. pardalinus* as an outgroup in (a) and *H. timareta*, *H. m. amaryllis, H. m. melponeme* collected from French Guiana, and *H. pardalinus* as an outgroup in (b). The value beside each reticulation edge (blue) represent inheritance probability.

#### 4.2.2 Ancient introgression in Primates

We selected a portion of the dataset of Vanderpool et al. (2020) that contains seven species in the Papionini group (three Asian species *Macaca nemestrina, M. fascicularis*, and *M. mulatta*; and four African species *Theropithecus gelada, Papio anubis, Cercocebus atys*, and *Madrillus leucophaeus*) where at least six introgression events were detected using the test statistic Δ, the method developed in Huson et al. (2005). We created an alignment of length 1,761,114 b.p. with eight sequences that includes the seven species listed above and an outgroup species *Callithrix jacchus*. We selected the outgroup species, *C. jacchus*, as it is appropriately distanced for our seven ingroup taxa based on the species tree provided in Vanderpool et al. (2020). We used the relationships presented in the species tree as our starting topology, and PhyNEST was executed following identical settings as in the *Heliconius* analyses with *h_max_* set to 0, 1, and 2. While the authors in the original study attempted to reconstruct a network with seven Papionini species with an outgroup using PhyloNet and SNaQ, they reported that both methods either failed to converge on an optimal network or yielded highly ambiguous results.

PhyNEST completed 10 independent runs for each *h_max_* using hill climbing and simulated annealing. When *h_max_* = 0, the tree topology was generally similar to the species tree presented in Vanderpool et al. (2020), although the four African species did not form a monophyletic clade. Figure 11 shows the estimated networks using *h_max_* = 1 and 2, both of which found using simulated annealing. When *h_max_* = 1, hybridization between *T. gelada* and *M. leucophaeus* was observed, with *γ* = 0.389 in the direction of *T. gelada*. When *h_max_* = 2, the same hybridization event between *T. gelada* and *M. leucophaeus* was observed, with *γ* = 0.378, in addition to the reticulation event among *Macaca* spp., *M. nemestrina* and *M. mulatta* with *γ* = 0.267 towards *M. mulatta*. While both reticulation events were not identified in Vanderpool et al. (2020) using Δ, the hybridizations identified in our networks are not unexpected, as approximately 40% of species in this group are known to hybridize (Tung and Barreiro, 2017). In fact, reticulation events identified in our analysis may be more probable than the scenarios proposed in Vanderpool et al. (2020), as our networks show hybridization events between members in the same continent, whereas the hybridizations detected in the original study were between species from different continents.

**Figure 11:**
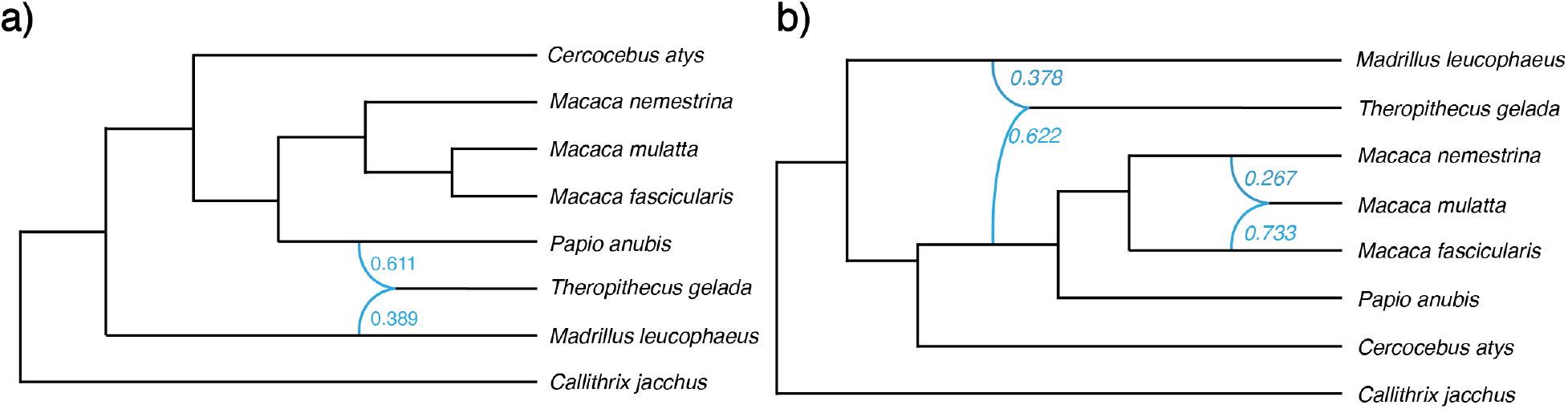
Phylogenetic networks of seven species in the Papionini group (plus an outgroup *Callithrix jacchus*) inferred with PhyNEST from the sequence alignment of 1,761,114 base pairs published in Vanderpool et al. (2020) using (a) *h_max_* = 1 and (b) =2. The value beside each reticulation edge (light and dark blue) represents inheritance probability for the edge with the same color.

## 5 Discussion

### 5.1 Significance and Limitations

The advances in next-generation sequencing technology over the last two decades have resulted in the unprecedented accumulation of large-scale genome data archived on public domains. While the expanded availability of genomic data undoubtedly provides opportunities for deeper understanding of the evolutionary histories among organisms, it simultaneously presents a series of challenges. For example, there is a shortage of efficient methods that extract phylogenetic signal from the data, use it to estimate relevant parameters, and measure statistical uncertainty in these estimates. The primary complication for developing such methods is that the set of potential histories that underlies the evolution of a given set of taxa is extremely large and exponentially increases with the number of taxa under consideration. This issue has come into sharper focus as the evidence of horizontal evolution (e.g., hybridization) in many species’ histories were found to be much more abundant than previously thought (Goulet et al., 2017; Grant and Grant, 1992; Soltis, 2013). A phylogenetic network model is now appreciated to be a better representation of many species’ evolutionary histories than a tree (Blair and Ané, 2020; Folk et al., 2018; Kong et al., 2022; Kong, 2022), however, the challenge of estimating the evolutionary history becomes more difficult as the dimensionality of the space of potential histories expands with reticulation.

Nevertheless, computational methods for inferring phylogenetic networks from genomic data are being actively developed under parsimony (Yu et al., 2013; Yan et al., 2022; Thomas et al., 2017), full (Yu et al., 2012, 2014) and composite likelihood (Yu and Nakhleh, 2015; Solís-Lemus and Ané, 2016, this study), Bayesian (Zhang et al., 2018; Flouri et al., 2020; Zhu et al., 2018; Wen et al., 2016), and other (Markin et al., 2022; Allman et al., 2019) frameworks. Among these, composite likelihood based methods are leading in overcoming scalability issues in network inference as they are shown to be more accurate than parsimony-based methods but more scalable than full likelihood methods (Solís-Lemus and Ané, 2016; Hejase and Liu, 2016). Unlike the full likelihood framework, the computational time for network estimation under the composite likelihood framework is generally independent of the number of loci or sequence length (Fig. 6) without compromising accuracy (Solís-Lemus and Ané, 2016).

PhyNEST is distinguished from the two existing network estimation methods that use a composite likelihood framework, namely SNaQ and PhyloNet, by the direct use of sequence data, rather than requiring the data to be summarized into a set of estimated gene trees prior to network inference. While such pre-processing of the data is a common strategy taken in the inference of both phylogenetic trees and networks using large multi-locus dataset to improve the speed of the analysis (e.g., ASTRAL (Mirarab et al., 2014), MP-EST (Liu et al., 2010), SNaQ (Solís-Lemus et al., 2017), PhyloNet (Than et al., 2008; Wen et al., 2018)), we argue that the overall computation time for estimating a network from the observed sequence data is not trivial because estimating a gene tree is still a computationally intensive process (Fig. 7). We show that the computation time for data pre-processing often exceeds the time required for the actual analysis, sometimes by orders of magnitude, when the number of loci is large. Parallelizing the gene tree estimation may mitigate this problem (Yin et al., 2019; Morel et al., 2019), but such computational capability may not be available to everyone.

The performance of the summarization approaches completely depends on the quality of the estimated gene trees, rather than the sequence data we observe (Molloy and Warnow, 2018; Roch and Warnow, 2015; Roch et al., 2019; Bayzid et al., 2015). Estimating an accurate gene tree can be hampered by a number of factors often beyond one’s control; for example, it is not uncommon for a dataset to contain closely related species, particularly when hybridization is suspected, and in this case, the signal in the sequence data might be too low to estimate accurate gene trees (Yu et al., 2014). Even when the sequence data contain enough signal as a whole, such signal can be diluted while summarizing it into gene trees. Estimating gene trees using the conventional phylogenetic tree inference methods often requires specification of nucleotide substitution models, and a misspecified model has been shown to bias phylogenetic tree estimation (Lemmon and Moriarty, 2004; Yang and Zhu, 2018; Swofford et al., 2001), although some have questioned the severity of this issue (Abadi et al., 2019). While our analysis assumed a correct substitution model when estimating gene trees from simulated data for SNaQ and PhyloNet to be conservative and maximize their performance, the effect of model misspecification on the accuracy and robustness of network estimation needs to be evaluated. In contrast, direct use of sequence data in PhyNEST is expected to fully utilize the phylogenetic signal available in data. While the current version implements the model introduced in Chifman and Kubatko (2015) that assumes the molecular clock, recent generalizations of the probability distribution of quartet site pattern frequencies to a relaxed clock as proposed in Richards and Kubatko (2022) could be implemented in PhyNEST in the future.

Another potential source of error in gene tree estimation is the misidentification of root position. Some summary-based methods (e.g., SNaQ) avoid this issue by allowing unrooted gene trees as input (and yielding a semi-directed network), but many methods require the input gene trees to be rooted. While rooting a topology using an outgroup is typically performed *ad hoc*, Mai et al. (2017) have shown high levels of rooting error due to gene tree–species tree discordance, which in turn can negatively influence those network methods that require rooted gene trees as input. Since the computation of site pattern probabilities implemented in PhyNEST involves rooted quartets, we also require a user to specify an outgroup taxon that will be used to root the network. This requirement can limit the use of PhyNEST in practice because a user must have *a priori* knowledge about the relationship between the taxa in their dataset, where the outgroup must be appropriately divergent from the ingroup so that it can be unambiguously segregated but does not result in long branch attraction simultaneously (Felsenstein, 1978). In a more practical sense, it is not always expected to have a genome-scale outgroup sequence readily available, particularly for non-model organisms. In order to overcome these problems, the rooting method developed in Tian and Kubatko (2017) that conducts a series of hypothesis tests for identifying a root position of a quartet tree using site pattern probabilities that can be extended to larger trees, seems to be an appropriate alternative that could be implemented in future versions of PhyNEST.

### 5.2 Future Directions for Scalable Network Estimation

The assessment of uncertainty of the clades in the estimated network is an essential step in phylogenetic analysis. In the tree context, this uncertainty can be measured using a bootstrap approach and a variety of methods, from nonparametric (Felsenstein, 1985; Efron, 1979) and parametric bootstrapping (Efron, 1985) to ultrafast (Minh et al., 2013) and little (Sharma and Kumar, 2021) bootstrap approaches, are available. Typically, a replicate alignment that has the same length as the original alignment is generated by resampling alignment columns from the original dataset, the tree from the replicate alignment is estimated (i.e., bootstrap replicate), and this procedure is repeated many times, generating a set of bootstrap replicates. The bootstrap replicates are either summarized into a single bootstrap consensus tree or the bootstrap value for a clade (i.e., the proportion of the replicate trees that recovered that particular clade) are mapped on the estimated tree on the tree that is estimated using the original alignment (see Janowitz et al. (2003) for a collection of excellent surveys on this topic).

For a set of bootstrap replicates, SNaQ implements the latter strategy by calculating the support for edges in the major tree (i.e., a parental tree obtained from a network by removing any reticulation edge with the inheritance probability < 0.5) then adds the support for the placement of each minor hybrid edges on that tree (Solís-Lemus and Ané, 2016). On the other hand, the full likelihood method implemented in PhyloNet employs a parametric bootstrap approach (Yu et al., 2014). Based on our current knowledge, no consensus method that combines a set of networks exists and a method to summarize bootstrap replicates into a single consensus network should be developed. Note this is different from the existing methods that aim to summarize a collection of conflicting trees into a single representative network (e.g. van Iersel et al., 2010; Huson and Rupp, 2008).

Since network space is very large and multi-dimensional, the network space in PhyNEST is arbitrarily shrunken using two strategies: first, the maximum number of dimensions of the network space is restricted by specifying *h_max_* by the user prior to the analysis; and second, it is assumed that the true network belongs to the class of level-1 TCTC phylogenetic networks for each *h_max_*. Although the network that represents the true evolutionary history may not always have the maximum number of reticulations that a network can have, it is possible to compute the upper bound of *h_max_*. Since the maximum number of vertices in a level-1 TCTC networks is expressed as |*V* (*N*)| ≤ 3*n* − 2 (van Iersel and Kelk, 2011) and |*V* (*N*)| ≤ 2*n* + 2*h* − 1 (Steel, 2016), where *n* and *h* represent the number of leaves and reticulation vertices, respectively, we can equate these two expressions and solve for *h* as 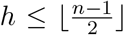. However, setting *h_max_* to this upper bound is not recommended since the likelihood favors more complex evolutionary relationships (i.e., more reticulations), and we expect the final network to have *h_max_* exactly. We stress that *h_max_* must be chosen carefully, since setting *h_max_* too low may miss the true network by not searching the space where it lies while setting *h_max_* too high will be computationally intensive and has potential to overestimate the amount of reticulation in the data. In SNaQ, it is suggested to estimate networks using various values of *h_max_* and conduct a model selection procedure to select the final network. In PhyloNet under the parsimony framework, improvement in the parsimony length is inspected as more reticulations are added, and stopped when the improvement is below a certain threshold (although a nonparametric bootstrap-based method to determine *h_max_* has been proposed in Park et al. (2010)).

In conclusion, estimating phylogenetic networks using the composite likelihood framework in PhyNEST is efficient and accurate, providing a method that scales up the inference of evolutionary history in the presence of hybridization to larger datasets. PhyNEST is distinguished from existing network inference methods by bypassing the data pre-processing step that summarizes sequence data into a set of gene trees. This maximizes the use of phylogenetic signal and prevents unwanted errors. Also, since the time required for gene tree estimation is not trivial, PhyNEST is expected to require less time for overall analysis. The estimated networks on two empirical dataset using PhyNEST demonstrate its practical applicability.

## 6 Supplementary Material

Data available from the Dryad Digital Repository: TBA

## 7 Acknowledgements

S.K. was supported by the Presidential Fellowship from the Ohio State University and Graduate Research Excellence Grant R. C. Lewontin Award by the Society for the Study of Evolution. The authors thank Jing Peng, Kristina Wicke, Cécile Ané, and Claudia Solís-Lemus for helpful advice in developing PhyNEST.

## 8 Appendix

## 8.1 Appendix A: Coefficients for Site Pattern Probabilities

**Table 1:**
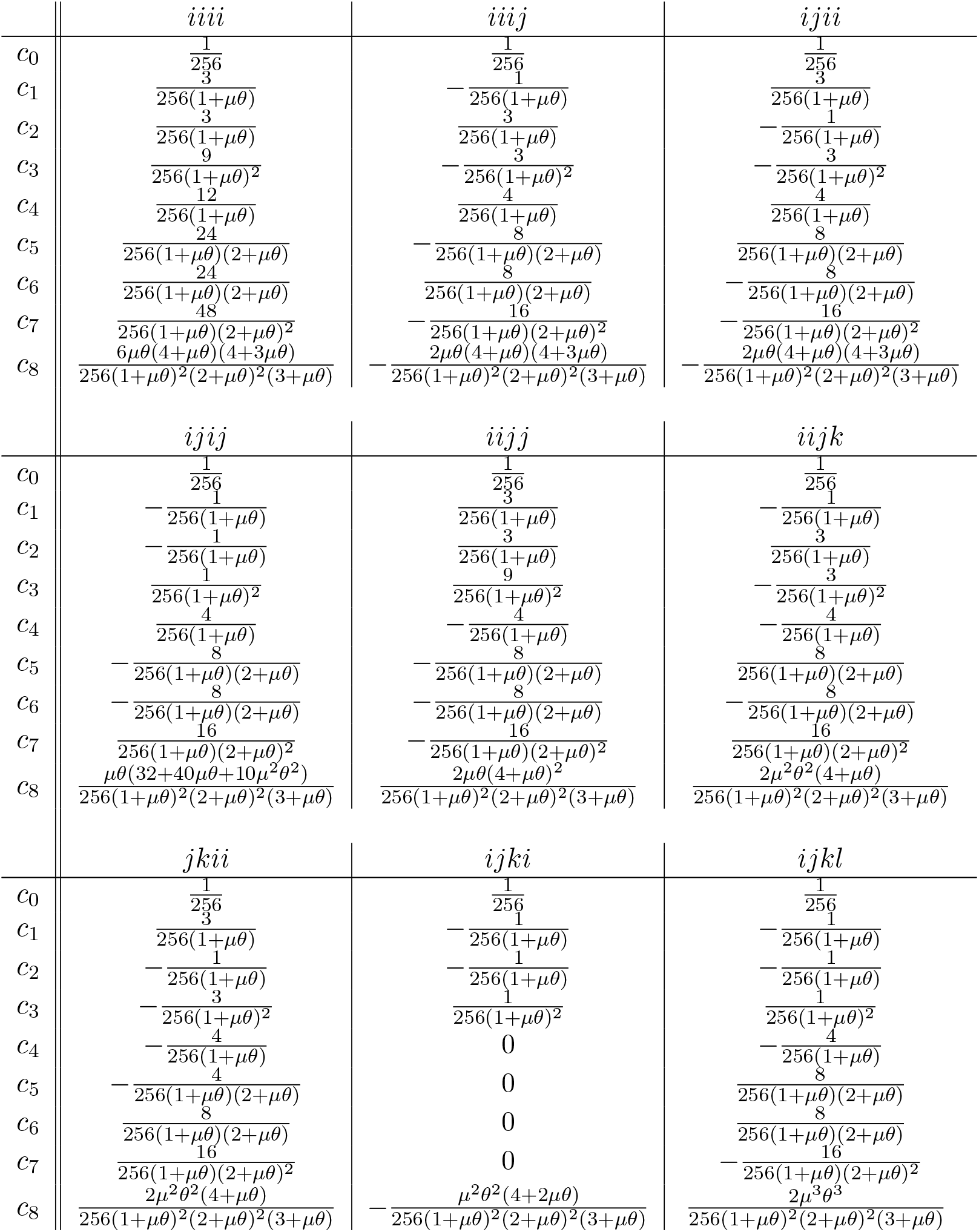
Coefficients for site pattern probabilities for rooted symmetric quartet tree

**Table 2:**
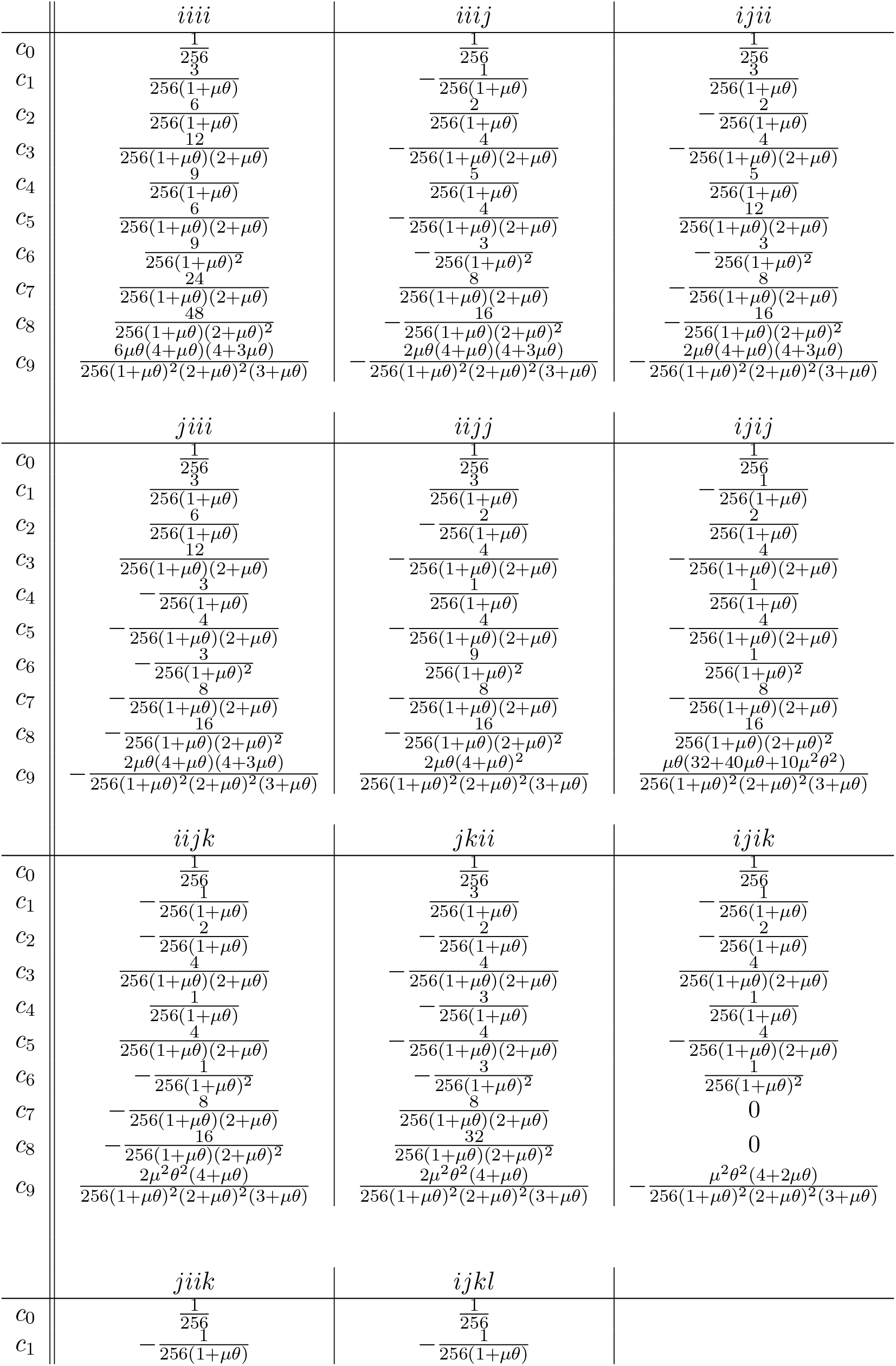
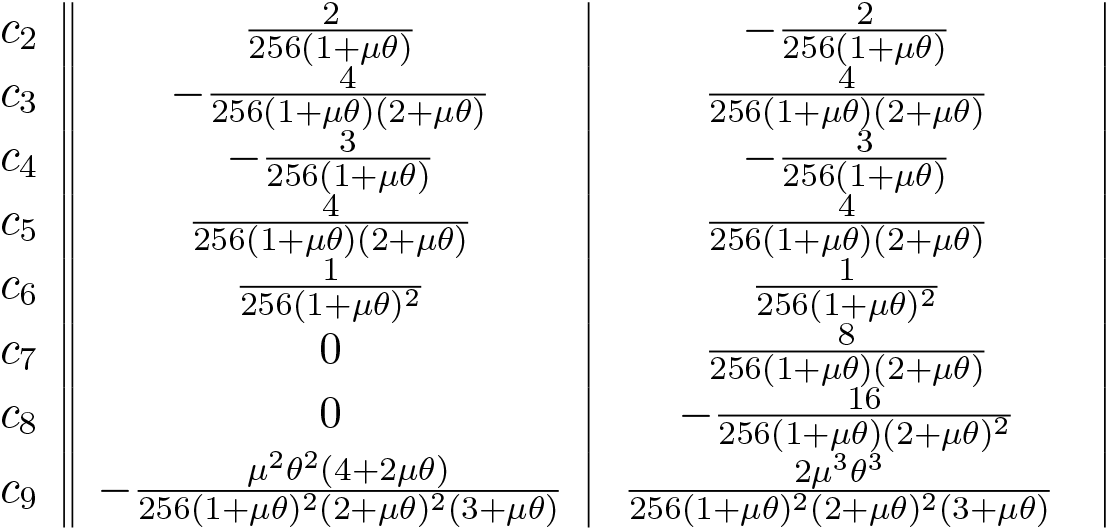
Coefficients for site pattern probabilities for rooted asymmetric quartet tree

## 8.2 Appendix B: The Relationships of the 15 Site Pattern Frequencies for 24 Permutations of Four Taxa

**Table 3:**
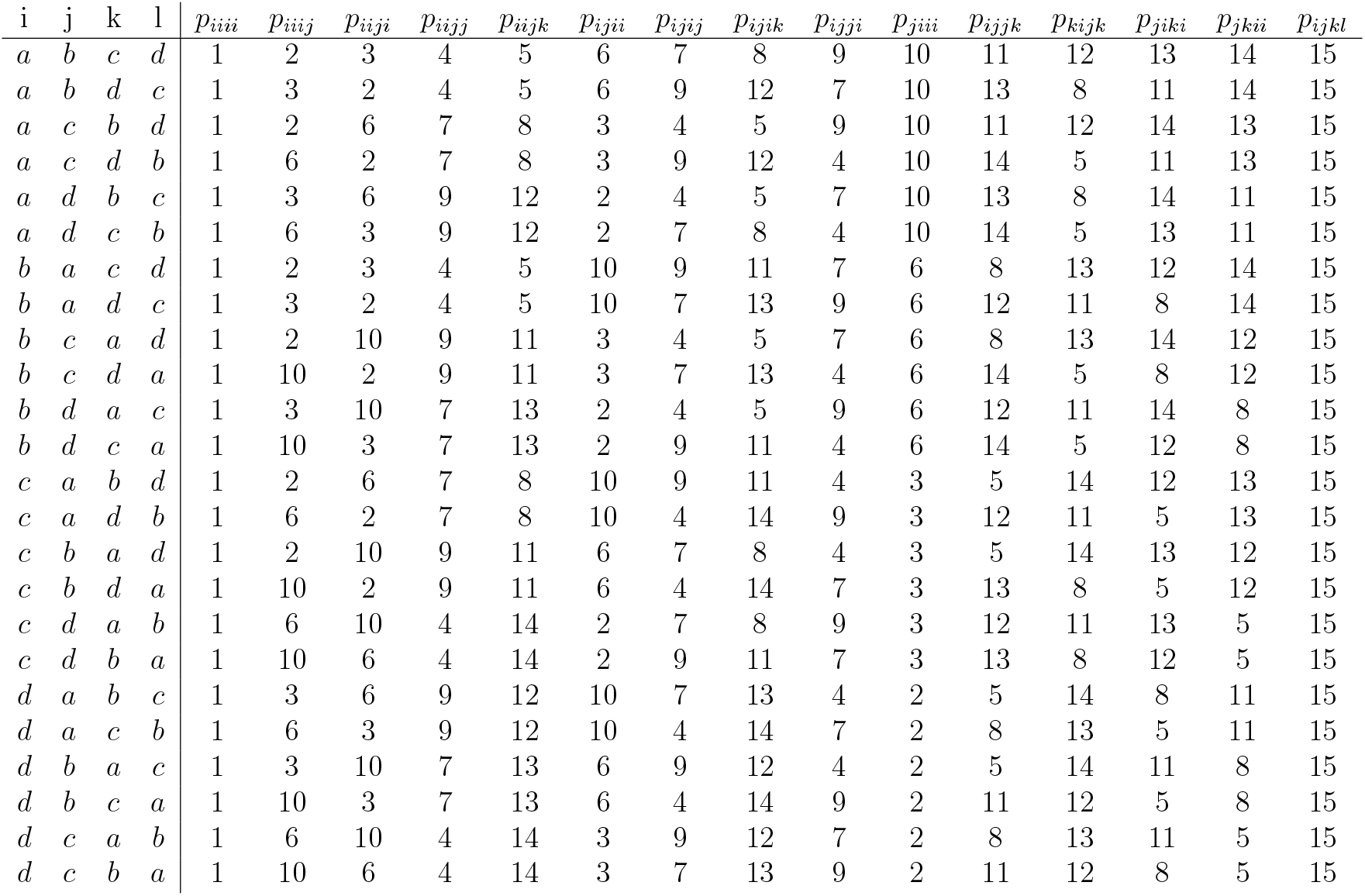
The relationships of the 15 site pattern frequencies for 24 permutations of four taxa *X* = {*a, b, c, d*}. The integers 1–15 are unique and this table illustrates that given the dataset, site pattern frequencies in all permutations of same set of four taxa are rearrangements of each other’s.

## 8.3 Appendix C: Computing Quartet Observed Site Pattern Probabilities

Let *W_q_* represent a vector of length 15 that contains site pattern counts for a quartet *q*, (*W_q_*)_*j*_ (see (10)) be an entry of the *W_q_* that represents the observed frequency of the site pattern *j*, *ω_j_* to be the weight of the site pattern *j* (see *ω* in (3)), and *M* to be the total length of the alignment,

## 8.3.1 Symmetric Quartet

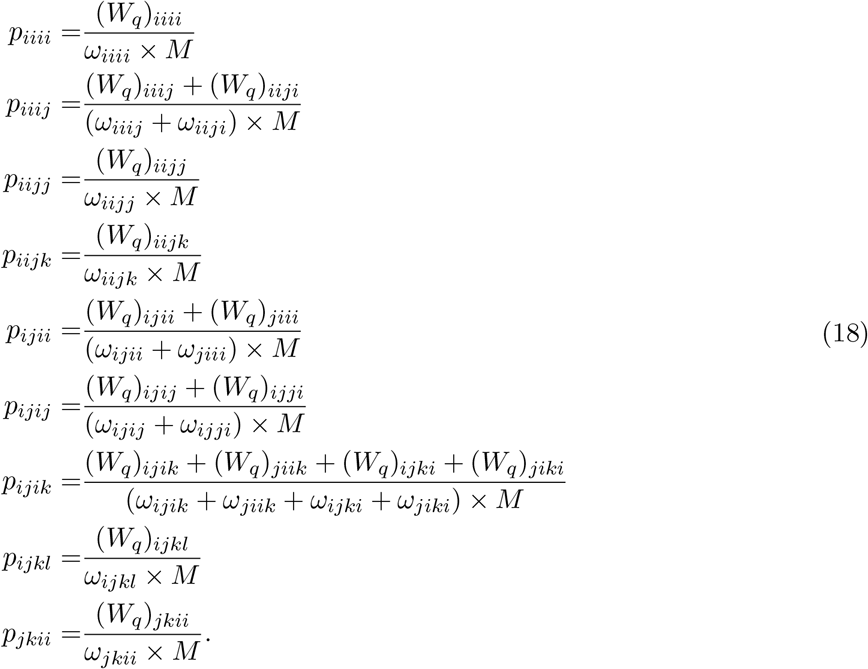

## 8.3.2 Asymmetric Quartet

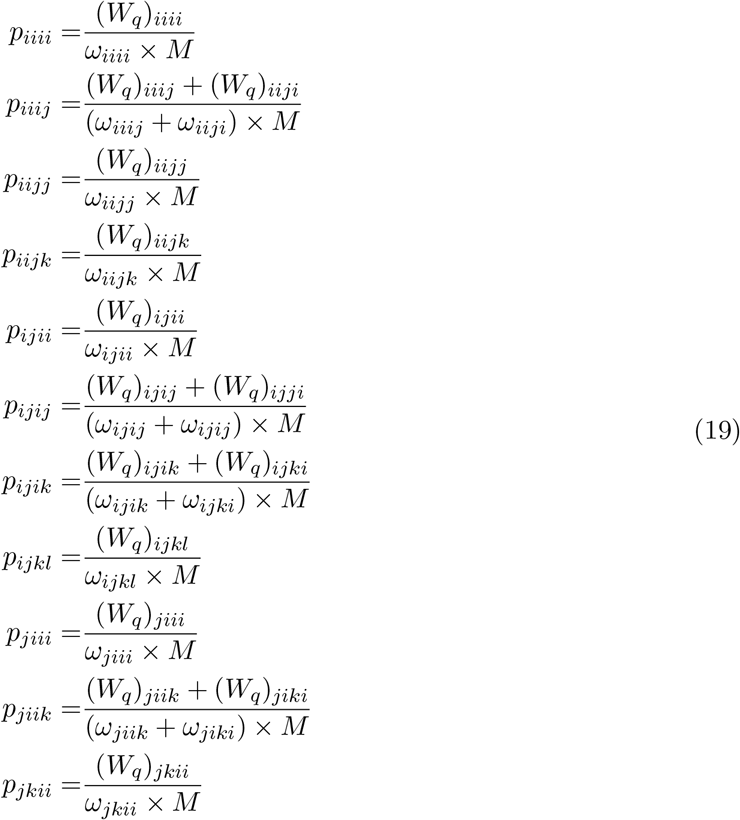

